# Microbiome-Derived Metabolites Shape CD4⁺ T-Cell Differentiation and Immune Aging in Chronic HIV-1 Infection

**DOI:** 10.64898/2026.01.13.699280

**Authors:** Amanda Cabral Da Silva, Luke Flantzer, Jaclyn Weinberg, Shuya Kyu, Lisa Daley-Bauer, Ana Carolina Santana, Aarthi Talla, Amber Lynn Rittgers, Sarah Welbourn, David Ezra Gordon, Jeffery Alan Tomalka, Vincent C. Marconi, Dean P. Jones, Souheil-Antoine Younes

## Abstract

The role of aromatic gut-derived bacterial metabolites (GDBMs) in shaping immune cell metabolism and function remains poorly explored. Using ex vivo metabolomic profiling of paired plasma and CD4⁺ T-cells from people living with HIV-1 (PLWH), we identified a network of aromatic GDBMs whose cell-associated abundance, rather than systemic levels, was linked to broad alterations in CD4⁺ T-cell metabolic and functional states. Among these metabolites, p-cresol sulfate (PCS) emerged as a mechanistic prototype investigated in depth. Ex vivo flow cytometry and single-cell RNA sequencing of CD4⁺ T-cells stratified by cell-associated PCS levels revealed dose-dependent enrichment of transcriptional programs associated with impaired differentiation capacity, regulatory-like identity, and cellular senescence. Consistently, in vitro transcriptomic and proteomic analyses of PCS-exposed CD4⁺ T cells demonstrated induction of cell-cycle arrest, mitochondrial dysfunction, and senescence-associated programs, including upregulation of p16 and p21. Integration of these immunometabolic features with measurements of HIV-1 reservoir size in PLWH revealed that CD4⁺ T-cell states defined by cell-associated GDBMs track with intact proviral DNA levels in vivo. Together, these findings define a microbiome-derived axis that reshapes CD4⁺ T-cell metabolism and fate and promotes immune aging–associated states in PLWH. Our data suggest that cell-associated GDBMs may foster immunometabolic CD4⁺ T-cell states previously linked to long-term HIV-1 reservoir persistence in vivo.

**Graphical Abstract.**
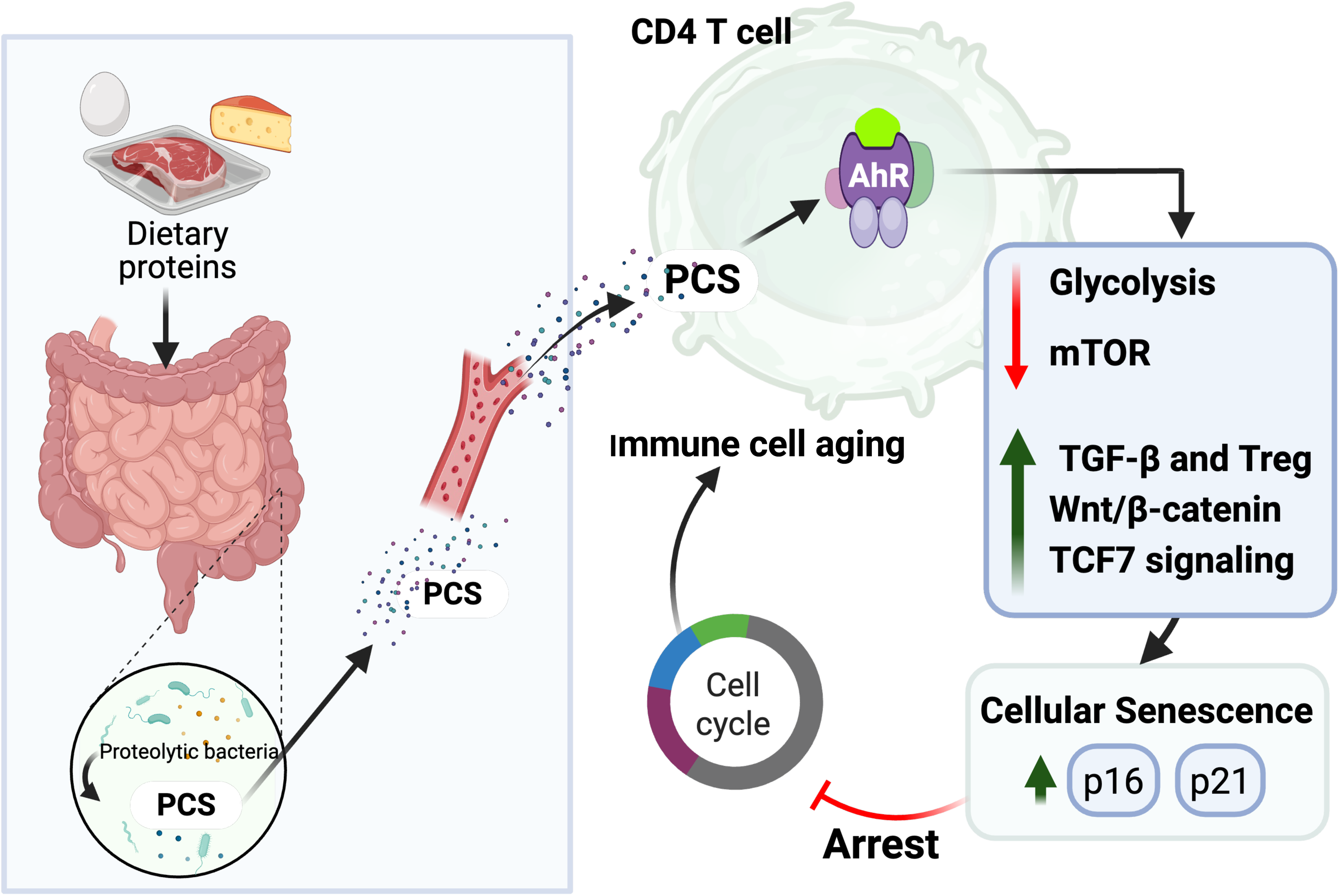
PCS-driven metabolic reprogramming and senescence promoting CD4+ T-cell immune cell aging. Dietary proteins are metabolized by proteolytic gut microbiota into p-cresol, which is absorbed and converted in the liver to PCS. Circulating PCS accumulates in CD4⁺ T-cells, where it activates the aryl hydrocarbon receptor (AhR). AhR signaling reduces glycolysis and mTOR activity, while enhancing TGF-β, Wnt/β-catenin, and TCF7 pathways, driving a regulatory-like and stem-like transcriptional program. These changes are associated with increased expression of p16 and p21, leading to cell cycle arrest and cellular senescence promoting CD4^+^ T-cell immune cell aging.

## Introduction

Gut-derived bacterial metabolites (GDBMs) are increasingly recognized as key modulators of host immunity and metabolic function. While short-chain fatty acids (SCFAs) (1–3) and microbiota-modified secondary bile acids (4, 5) are well characterized for their roles in immunity, less is known about the effects of other GDBMs, particularly aromatic metabolites such as p-cresol sulfate (PCS), p-cresol glucuronide (PCG), phenylacetylglycine (PAG), and indole-3-acetic acid (IAA), which are generated through bacterial degradation of amino acids by the gut bacterial flora(6). These metabolites can reach peripheral tissues to exert immunomodulatory effects, but their functional impact on human T-cells remains largely unexplored. Among these, PCS, a prototypical GDBM, accumulates in circulation during conditions of dysbiosis, chronic inflammation, and renal dysfunction(6–9). PCS originates from bacterial degradation of tyrosine and phenylalanine in the gut, is sulfated in the liver, and normally cleared by the kidney (7). Elevated PCS levels is associated with oxidative stress(10), endothelial dysfunction(11), and impaired immune responses in chronic disease states (12, 13). However, the cell-intrinsic effects of PCS on adaptive immunity, particularly CD4⁺ T-cell function and fate, remain poorly defined. PCS is proinflammatory and induces the production of leukocyte free radicals and the release of endothelial microparticles (11). PCS impairs mitochondrial dynamics and functions of renal tubular cells (14) as well as in leukocytes (15). We have investigated the role of PCS in people living with HIV-1 (PLWH), particularly focusing on immune nonresponders (INRs) who, despite effective antiretroviral therapy (ART), fail to restore normal CD4⁺ T-cell counts (16). The study revealed that INRs exhibited elevated plasma and CD4⁺ T-cell levels of PCS. Plasma PCS levels negatively correlated with CD4⁺ T-cell counts, suggesting a potential mechanistic link between PCS accumulation and impaired immune recovery. Further, in vitro analyses demonstrated that exposure of healthy CD4⁺ T-cells to PCS led to several detrimental effects: 1) Inhibition of proliferation: PCS exposure resulted in a significant reduction in CD4⁺ T-cell proliferation. 2)

Induction of apoptosis signaling pathway: there was an increase in programmed cell death signaling among these T cells upon PCS treatment, yet the levels of Annexin V detected did not reach more that 20% at the highest in vitro PCS concentration. 3) Mitochondrial dysfunction: PCS diminished the expression of essential mitochondrial proteins, leading to observable perturbations in mitochondrial networks, as evidenced by electron microscopy (8).

CD4⁺ T-cells are central orchestrators of immune responses, balancing effector cytokine production with long-term memory and regulatory functions. Their activation and differentiation are tightly coupled to metabolic reprogramming, with key pathways such as mTOR, glycolysis, and oxidative phosphorylation supporting lineage commitment and effector function (17–19). Disruption of these metabolic circuits can compromise T-cell fate decisions, promote dysfunction, or lead to the emergence of stem-like or exhausted states, particularly under conditions of chronic antigen exposure or metabolic stress(20). Senescence in CD4⁺ T-cells, characterized by cell cycle arrest, loss of proliferative capacity, and altered metabolic programs, is increasingly recognized as a feature of both immunological aging and chronic infections (21–23). This phenotype is associated with mitochondrial dysfunction, altered redox balance, and the expression of canonical regulators such as p16 and p21 proteins (21, 24).

HIV-1 infection accelerates immune cell aging through multiple mechanisms, many of which mirror natural immunosenescence observed in elderly individuals (25, 26). Chronic immune activation is a hallmark of HIV infection, driving repeated cycles of immune cell proliferation, activation, and turnover that ultimately exhaust regenerative capacity (27, 28). CD4⁺ T-cells undergo replicative senescence, reduced proliferative potential, increased expression of senescence markers such as p16 (29). Together, these processes compromise immune homeostasis and diminish the capacity to respond to new antigens, contributing to the premature immune aging observed in PLWH (30–33).

CD4⁺ T-cells are the primary cellular reservoir for latent virus, with long-lived memory cells enabling persistent viral latency despite ART (34). Understanding the metabolic constraints that maintain these cells is critical. Several studies have shown that metabolic features of CD4⁺ T-cells correlated with reservoir stability (35–39) yet the upstream triggers remain poorly defined. Recent studies have shed light on the metabolic underpinnings of HIV-1 persistence in CD4^+^ T-cells, offering new insights into reservoir biology and potential therapeutic targets. Valle-Casuso et al.(40) demonstrated that CD4⁺ T-cells with elevated glycolytic and oxidative phosphorylation activity are more susceptible to HIV-1 infection, and that metabolic inhibition of these pathways suppress viral replication and selectively eliminate infected cells, positioning metabolism as a determinant of reservoir seeding. Similarly, Akiso et al. (41) observed metabolic reprogramming in CD4⁺ T-cells from virologically suppressed individuals on ART, showing increased mitochondrial membrane potential and mass but reduced glucose and fatty acid uptake, which may support cell survival and latency. Kang and Tang (42) further highlighted the role of the PI3K/Akt/mTOR axis in upregulating glucose metabolism during infection, thereby facilitating viral replication and potentially supporting latency through anabolic signaling. Expanding on this, Taylor et al. (43) provided mechanistic evidence that mTOR activation overcomes metabolic blocks to enable efficient HIV reverse transcription and nuclear import, linking mitochondrial and nutrient signaling directly to early infection events. Lastly, Wan et al. (44) used multi-omics profiling to associate metabolic dysregulation with poor immune recovery despite viral suppression, indicating that alterations in lipid and amino acid metabolism may impact reservoir size and T cell functionality. Despite these advances, the upstream metabolic triggers that shape reservoir-permissive CD4⁺ T-cell states remain largely undefined.

The impact of GDBMs on the metabolic programming and fate decisions of CD4⁺ T-cells in HIV-1 infection settings, including immune aging, differentiation, senescence, and functional capacity, remains largely unexplored. Here, we combine multi-omics profiling, including transcriptomics, proteomics, cytokine analysis, and metabolomics, with functional in vitro assays and ex vivo validation in PLWH to dissect the effects of PCS on CD4⁺ T-cell biology. We demonstrated that PCS exposure drives CD4⁺ T-cell senescence and promotes a regulatory-like transcriptional program, while concurrently restraining effector differentiation. PCS represses mTOR and effector cytokine programs while activating TGF-β, the aryl hydrocarbon receptor (AhR), and Wnt signaling, and reshapes the T cell proteome toward mitochondrial stress and metabolic adaptation. PCS exemplifies a microbial metabolite that reshapes CD4⁺ T-cell fate through metabolic and transcriptional reprogramming and may contribute to an immune environment conducive to HIV-1 reservoir persistence.

## Results

### Activation-Dependent Cell Association of PCS in CD4⁺ T-Cells Is Independent of Plasma Levels

To determine whether PCS levels in plasma reflect cell association in CD4⁺ T-cells, we quantified PCS the concentrations in matched plasma and sorted CD4⁺ T-cells from 50 PLWH. Mass spectrometry analysis revealed that cell-associated PCS levels were significantly lower than those in plasma, with mean concentrations of 0.025 μM in CD4⁺ T-cells versus 11.8 μM in plasma (*P* < 0.0001) (**Supplementary Figure 1A**). Despite this large disparity, no significant correlation was observed between plasma and cell-associated PCS levels (Spearman r = –0.1861, P = 0.1957) (**Supplementary Figure 1B**), suggesting that circulating PCS levels do not reliably predict CD4^+^ T-cell association. This may be due, in part, to the fact that approximately 95% of PCS in plasma is tightly bound to albumin (7), leaving only a small free fraction available for cellular entry. Additionally, PCS may engage tissues and immune cell subsets other than CD4⁺ T-cells, further decoupling systemic concentrations from cell-associated levels in CD4⁺ T-cells. To further investigate the kinetics and determinants of PCS cell-association, CD4⁺ T-cells from healthy donors were incubated with 50 or 100 μM PCS for 1, 3, or 6 days. PCS uptake was detectable at all time points and was dose-dependent, with cell-association concentrations ranging from 4 to 8 nM at 50 μM PCS and ∼8–16 nM at 100 μM PCS (**Supplementary Figure 1C**). Notably, T-cell receptor (TCR) stimulation with anti-CD3/CD28 during PCS exposure on day 6 enhanced PCS cell-association. Together, these data demonstrate that PCS associates with CD4⁺ T-cells in a dose- and activation-dependent manner, independent of circulating protein-bound PCS concentrations, which fail to predict CD4⁺ T cell–associated PCS levels. Importantly, exposure to 100 μM PCS results in quantifiable nanogram-scale levels of cell-associated PCS in CD4⁺ T-cells.

### Cell-Associated GDBMs in CD4⁺ T Cells from PLWH Associate with TCA Cycle Disruption and Global Metabolic Dysregulation

We then performed untargeted metabolomics along with cell-associated PCS, PAG, PCG, and IAA quantification by targeted metabolic analysis. Metabolic profiling was performed on lysate of CD4⁺ T-cells from 26 immune-responders PLWH (listed below **Figure 1A**, and **1B**). As shown in **Figure 1A**, we observed a gradient of cell-associated PCS concentrations across individuals that significantly stratified the abundance of multiple tricarboxylic acid (TCA) cycle intermediates. Higher cell-associated PCS levels were associated with elevated levels of key metabolites such as citrate, α-ketoglutarate, succinate, malate, and fumarate, suggesting accumulation of incompletely oxidized TCA intermediates consistent with mitochondrial dysfunction. In contrast, plasma PCS concentrations did not show the same degree of association with TCA metabolites (**Figure 1B**), indicating that cell-associated PCS levels, not circulating levels, are more reflective of T-cell metabolic perturbation. Then, Spearman correlation analysis (**Figure 1C**) confirmed that PCS levels within CD4⁺ T-cells were positively correlated with several central TCA metabolites, including AMP, succinate, and fumarate, further supporting PCS-associated bioenergetic stress and mitochondrial inefficiency. To extend beyond TCA cycle alterations, we used untargeted metabolomics to assess how PCS cell-level and related GDBMs (PAG, PCG, and IAA) modulate broader metabolic pathway activity in CD4⁺ T-cells. In **Figure 1D**, cell-level PCS showed the broadest and most coherent associations with CD4⁺ T-cell metabolism. Higher CD4⁺ T-cell-associated PCS concentrations positively correlated with multiple core pathways, most prominently central carbon/energy and biosynthetic programs including the TCA cycle, glycolysis/gluconeogenesis, purine and pyrimidine metabolism, and several amino-acid catabolic routes. PCS also correlated with pathways related to redox balance and cofactor metabolism, consistent with broad metabolic remodeling linked to PCS accumulation. Importantly, these pathway-level correlations were detected for cell-associated PCS, reinforcing that cellular retention, not plasma abundance, aligns with metabolic remodeling. Expanding beyond PCS, PAG displayed a correlation pattern that largely mirrored PCS. Cell-associated PAG showed a similar metabolic footprint to PCS, positively correlating with pathways related to energy production, nucleotide biosynthesis, and amino acid processing. This similarity suggests that PCS and PAG participate in a shared aromatic GDBM-associated metabolic network within CD4⁺ T-cells, with both metabolites tracking along the same direction of pathway perturbation. By contrast, PCG showed an opposing relationship to this PCS/PAG axis. Cell-associated PCG concentrations correlate negatively with several of the same pathways that rose with PCS and PAG, including energy-linked and biosynthetic programs. These inverse associations position PCG within the same aromatic GDBM family yet suggest a metabolically divergent role in CD4⁺ T-cell regulation, implying heterogeneity in how structurally related GDBMs map onto CD4⁺ T-cell metabolic states. Finally, IAA did not display significant associations with CD4⁺ T-cell metabolic pathways in this dataset.

**Figure 1.**
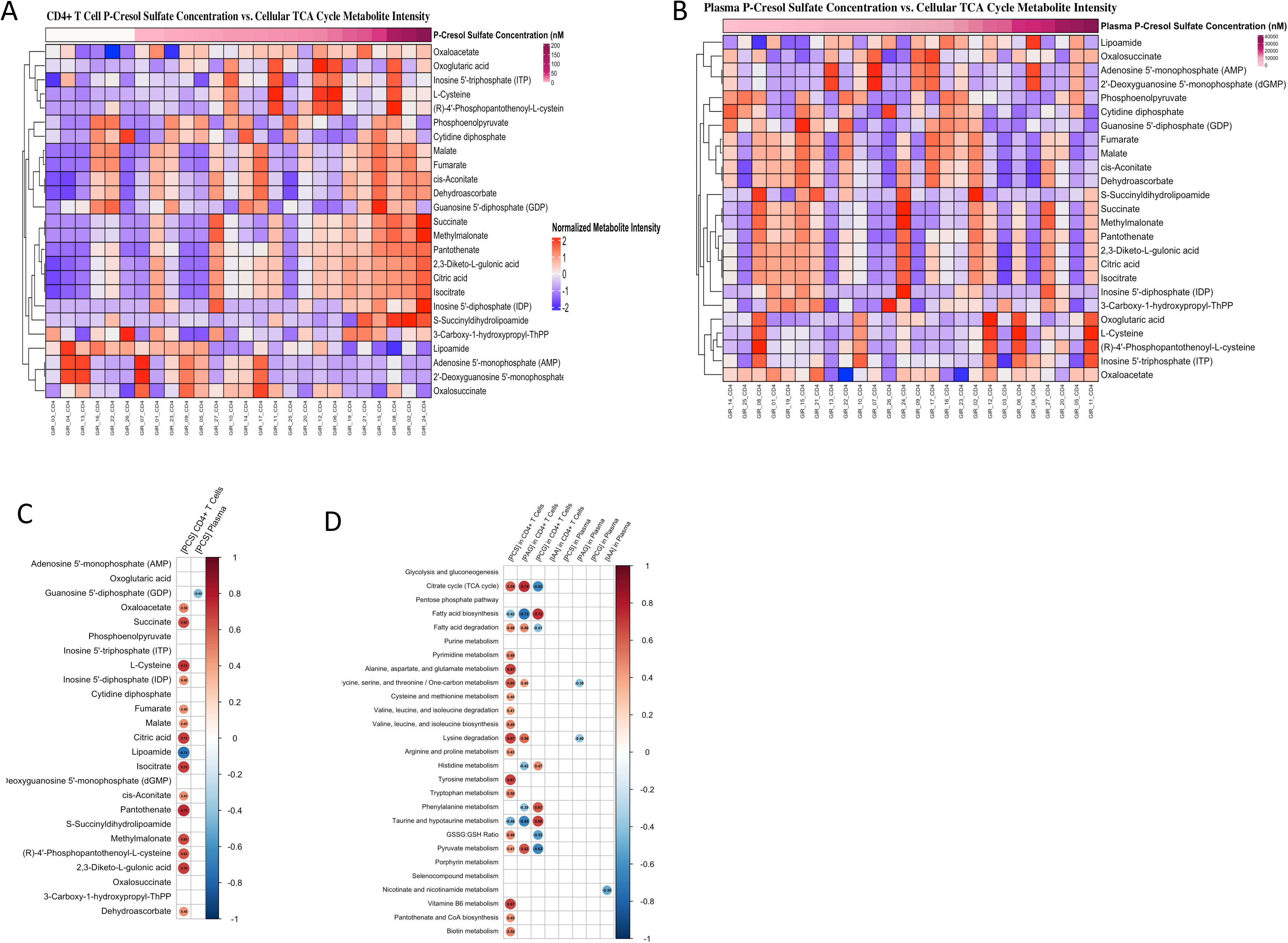
Metabolite Profiling and PCS Quantification in CD4⁺ T-Cells from PLWH. CD4⁺ T-cells were sorted from PBMCs of 26 immune responders PLWH. Cell-associated and plasma concentrations PCS were quantified by targeted mass spectrometry. Untargeted metabolomics analysis was performed on sorted CD4⁺ T-cells using high-resolution LC-MS. (**A**) and (**B**) display normalized intensities of tricarboxylic acid (TCA) cycle metabolites stratified by cell-associated PCS (**A**) or plasma PCS (**B**) concentrations. (**C**) Spearman correlation coefficients were calculated between cell-associated PCS levels and individual TCA metabolites. (**D**) Spearman correlation heatmap showing associations between the concentrations of PCS, PAG, PCG, and IAA measured in CD4⁺ T-cells or plasma, and the enrichment of metabolic pathways identified by untargeted metabolomics. Rows represent metabolic pathways, and columns represent cell-associated or plasma metabolite levels. Circle size and color reflect the magnitude and direction of the correlation (red: positive; blue: negative).

Together, these findings demonstrate that cell-associated accumulation of specific aromatic GDBMs, particularly PCS and PAG, is tightly linked to global metabolic reprogramming in CD4⁺ T-cells from PLWH. PCS exhibited the strongest and most consistent associations with disrupted energy metabolism and biosynthetic pathways, reinforcing its role as a key immunometabolic effector. Of the 27 metabolic pathways assessed, cell-associated PCS concentrations correlated with 19 pathways, reflecting the broadest metabolic impact among the GDBMs analyzed. PAG correlated with 9 pathways, displaying partial overlap with PCS, while PCG was associated with 8 pathways, often in an opposing direction to PCS and PAG. Importantly, these metabolic alterations were specific to cell-associated GDBM abundance and were not reflected by plasma concentrations, underscoring the importance of considering cellular association when assessing the immunometabolic impact of GDBMs.

### Cell-associated GDBMs correlate with metabolic pathways linked to the intact HIV-1 reservoir

Long-term persistence of the HIV-1 reservoir is tightly linked to CD4⁺ T-cell homeostasis, survival, and differentiation (45). While previous studies have implicated immune activation and cellular metabolism in reservoir maintenance (40, 42, 46), direct assessments of how cell-associated metabolic states influence reservoir size remain limited. As our study leveraged untargeted metabolomic profiling of primary CD4⁺ T-cells, we sought to determine whether metabolic pathway activity within the CD4⁺ T-cell compartment associates with HIV-1 reservoir metrics. In parallel, given the strong linkage between cell-associated GDBMs and CD4⁺ T-cell metabolic remodeling, we further examined whether cell-associated GDBM abundance tracks with metabolic pathways implicated in reservoir persistence. To address this, we examined whether the metabolic pathways associated with GDBMs cell-level are linked to HIV-1 reservoir persistence, as measured by the intact proviral DNA assay (IPDA). Global correlation analysis between CD4⁺ T-cell metabolite abundances and intact HIV-1 reservoir size revealed a bifurcated pattern of associations (**Figure 2A**). Of the 342 detected cell-associated metabolites, 92 showed significant correlations with intact proviral DNA, segregating into metabolites negatively or positively associated with reservoir size. **Figure 2B** illustrates the strong positive correlation between intact and total HIV-1 DNA measured in matched CD4⁺ T-cell samples, validating the reservoir quantification and enabling comparative metabolic analyses. To resolve pathway-level relationships, we next examined the top metabolites contributing to negative (**Figure 2C**) or positive (**Figure 2D**) correlations with intact reservoir size. Metabolites negatively associated with the intact reservoir were enriched in pathways related to nucleotide biosynthesis, amino acid metabolism, and mitochondrial function, whereas metabolites positively associated with the intact reservoir mapped to pathways involved in lipid metabolism, purine and pyrimidine metabolism, and the tricarboxylic acid cycle. Notably, individual GDBMs, including PCS, PAG, PCG, and IAA, did not directly correlate with intact proviral DNA; however, PCS and PAG were positively associated with metabolic pathways that were themselves positively correlated with intact reservoir size, whereas PCG and IAA were linked to pathways exhibiting negative associations (**Figure 2C–D**).These divergent patterns suggest that GDBMs collectively influence the immunometabolic landscape of CD4⁺ T cells rather than acting as direct correlates of reservoir burden. Together, these data identify cell-associated GDBMs as previously unrecognized contributors to metabolic programs associated with HIV-1 reservoir persistence and latency in vivo.

**Figure 2.**
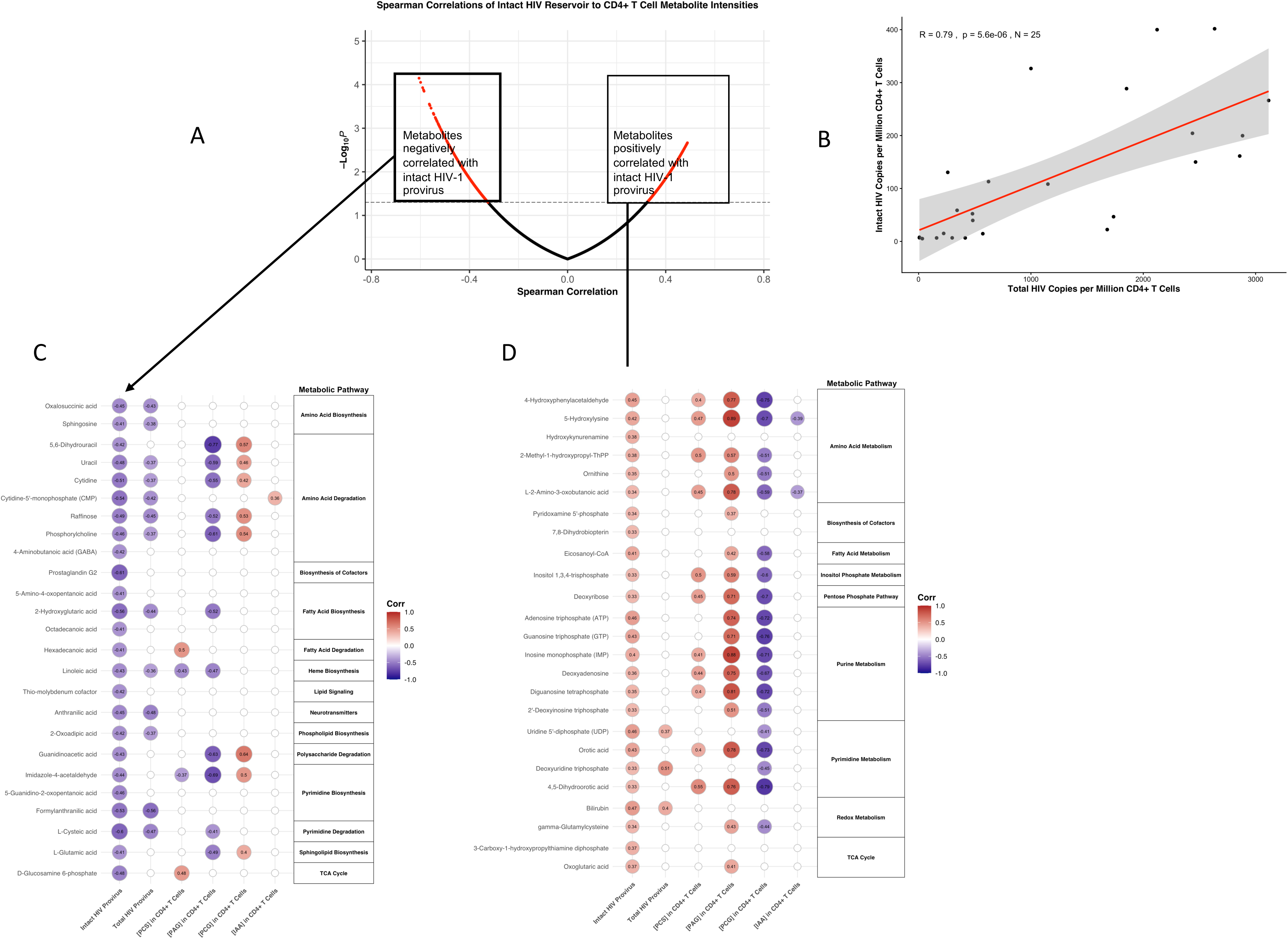
Metabolic pathway correlations with intact and total HIV-1 reservoir levels in CD4⁺ T-cells and relationship to the GDBMs. (**A**) Volcano plot showing Spearman correlations between CD4⁺ T-cell metabolite intensities and intact HIV-1 reservoir size. Red points indicate statistically significant correlations (p < 0.05), and boxed regions highlight metabolites with strong negative (left) and positive (right) associations. (**B**) Linear regression of intact vs. total HIV-1 copies per million CD4⁺ T cells, demonstrating a strong positive correlation (R = 0.79, p = 5.6e-06, n = 25). (**C**) Heatmap of negatively correlated metabolites (highlighted in panel A, left box), categorized by metabolic pathways. (**D**) Heatmap of positively correlated metabolites (highlighted in panel A, right box), with pathway annotations. Circle size and color scale represent strength and direction of Spearman correlation coefficients between each metabolite and reservoir size (intact and total IPDA values). Targeted quantification was performed for PCS, PAG, PCG, and IAA. Values represent Spearman correlation coefficients between the indicated GDBMs (x-axis) and metabolites within key pathways (y-axis). Pathway annotations on the right summarize functional categories significantly associated with GDBM levels. Positive correlations (red) indicate co-enrichment between GDBMs and metabolites, while negative correlations (blue) indicate reciprocal relationships suggestive of metabolic suppression. Data were obtained ex vivo from freshly isolated CD4⁺ T-cells (n = 26 donors).

### Cell-Associated PCS Levels Are Linked to Altered Marker Expression and Shifts in CD4⁺ T-Cell Subsets Ex vivo

We then focused our analysis on PCS to determine how cell-associated PCS concentrations shape CD4⁺ T-cell biology ex vivo. We stratified peripheral blood samples from PLWH into four groups based on cell-associated PCS concentrations measured in CD4⁺ T-cells: no PCS, low PCS, medium PCS, and high PCS, with six individuals per group as listed in **Figure 3A**. This stratification enabled us to assess how PCS cell-level correlates with CD4⁺ T-cell phenotype and subset composition ex vivo without external stimulation. Using concatenated FlowJo files for each group, we analyzed the median fluorescence intensity (MFI) of key surface and intracellular markers associated with differentiation, homeostasis, proliferation, and regulatory function. A clear reduction in CD4 expression was observed with increasing PCS levels (**Figure 3B**). Concurrently, there was a progressive increase in the expression of TCF7, CCR7, CD45RA, FOXP3, and CD25, all of which are associated with naïve-like, central memory (TCM), or regulatory T cell (Tregs) programs. In contrast, markers of proliferation and activation, namely Ki-67 and CD71, exhibited a notable decrease with rising PCS levels, reinforcing the interpretation that high PCS may impair CD4⁺ T-cell proliferative capacity. We then applied t-distributed Stochastic Neighbor Embedding (t-SNE) visualization flow self-organizing map (FlowSOM) to assess CD4⁺ T-cell landscape changes across PCS concentration groups (**Figure 3C** and **Supplementary Figure 2**). With increasing PCS, a visible shift in subset distribution emerged. Specifically, there was a marked reduction in the terminally differentiated effector memory CD45RA (TEMRA) or cytotoxic CD4^+^ T-cell population; in contrast, Tregs FOXP3⁺ showed a significant increase in frequency across the PCS gradient, including Ki-67⁺ proliferative Tregs (**Figure 3C**). Quantitative analysis confirmed these patterns. TEMRA cells significantly declined, while Tregs (total and Ki-67⁺ Tregs) exhibited a statistically significant increase in frequency at higher PCS concentrations (**Figure 3D**). Although not statistically significant, there was a consistent trend toward increased frequencies of naïve, TCM, and TEM (effector memory) CD4⁺ T-cells in the mid and high PCS groups. Additionally, a trend toward decreased Ki-67⁺ TEM cells was observed, further supporting a PCS-associated reduction in CD4⁺ T-cell proliferative activity.

**Figure 3.**
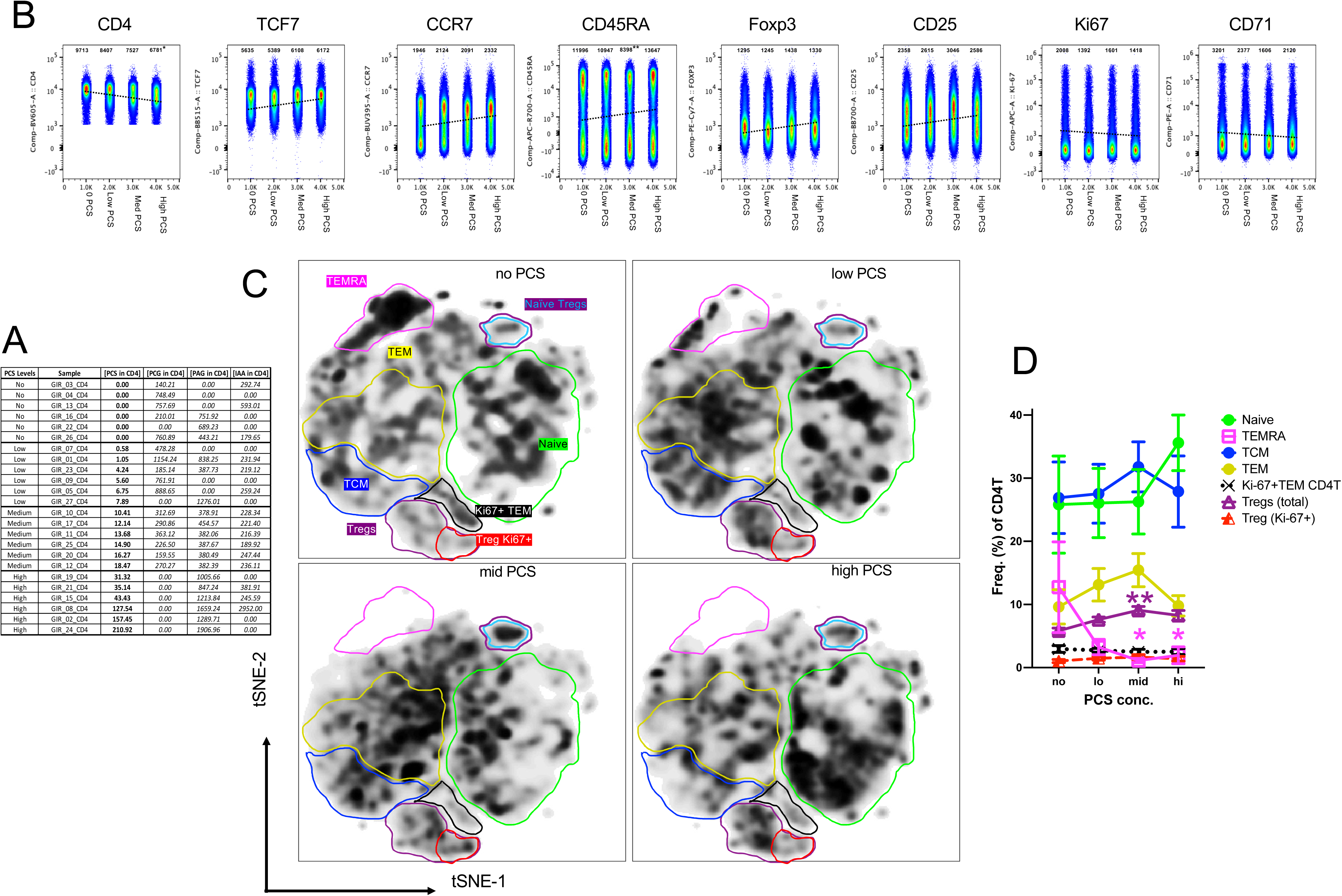
Ex vivo phenotypic characterization of CD4⁺ T-cells stratified by cell-associated PCS levels in PLWH. **(A**) Table summarizing cell-associated PCS concentrations in CD4⁺ T-cells across individual donors (n = 24), grouped into four categories based on PCS levels: no PCS, low PCS, medium PCS, and high PCS (6 donors per group). Concentrations (nM) are shown in bold for PCS, and in italics for PAG, PCG, and IAA in each sample. (**B**) Flow cytometric pseudoplots showing CD4, TCF7, CCR7, CD45RA, FOXP3, CD25, Ki-67, and CD71 expression. Plots represent concatenated files for each PCS group. Median fluorescence intensity (MFI) values for each marker are indicated numerically within individual plots. (**C**) t-SNE plots of CD4⁺ T-cells demarked by FlowSOM-defined clusters corresponding to canonical T cell subsets, including naïve, TCM, TEM, TEMRA, total Tregs, naïve Tregs and Ki-67⁺ Tregs. Plots represent pooled data from each PCS group. (**D**) Quantification of CD4⁺ T-cell subset frequencies as a percentage of total CD4⁺ T-cells in each PCS group. Data represent mean ± SD. Statistical significance was assessed using one-way ANOVA with multiple comparisons correction. *p<0.5, **, p<0.01.

Together, these immunophenotypic changes parallel the metabolic alterations described in **Figure 1**, where higher cell-associated PCS concentrations correlated with disruptions in TCA cycle intermediates and multiple metabolic pathways. The observed reduction in Ki-67 and CD71, coupled with the accumulation of TCF7⁺, CCR7⁺, and FOXP3⁺ cells, may reflect a metabolically constrained environment, likely shaped by PCS-induced mitochondrial dysfunction or impaired bioenergetics. Taken together with the metabolic alterations observed in **Figure 1**, these data suggest that cell-associated PCS modulates CD4⁺ T-cell metabolism and differentiation, promoting a low-proliferative, Treg-skewed phenotype with reduced effector turnover.

### Single-cell transcriptomics reveals that cell-associated PCS skews CD4⁺ T-cells toward regulatory, stem-like, and senescence-associated transcriptional states ex vivo

To investigate whether cell-associated PCS concentrations shape the transcriptional landscape of CD4⁺ T-cells ex vivo, we performed scRNA-seq on sorted CD4⁺ T-cells from six participants: three donors with high PCS (GIR-24, GIR-08, GIR-02) and three with low PCS levels (GIR-07, GIR-03, GIR-04). This analysis aimed to validate and expand upon the phenotypic differences previously observed using flow cytometry and FlowSOM clustering. As shown in **Figure 4A**, UMAP projections revealed clear density differences between low and high PCS samples, suggesting PCS-associated shifts sub-population distribution. Clustering analysis of all CD4⁺ T-cells identified 12 transcriptionally distinct clusters (**Figure 4B**), which were annotated into known CD4⁺ T-cell subsets (naïve, TCM, TEM, CTL, and proliferating) using established classification algorithms (**Figure 4C**). Differential abundance analysis (**Figure 4D**) showed that clusters C5 and C0 were significantly enriched in cells from high PCS donors, while clusters C2 and C6 were more abundant in low PCS donors. Annotation of the cluster compositions revealed that clusters C5 and C0 were predominantly composed of Tregs and TCM, respectively, whereas C2 and C6 were enriched for TEMRA-like CD4⁺ cytotoxic T lymphocytes (CTL) and TEM cells (**Figure 4E**). Importantly, these patterns were consistent across individual donors as shown in **Figure 4F**, underscoring the reproducibility of PCS-associated transcriptional states. To further explore the molecular signatures associated with PCS levels, we performed gene expression comparisons between clusters enriched in high vs. low PCS donors. As shown in **Figure 4G**, CD4⁺ T-cells from high PCS participants upregulated gene modules associated with the AHR pathway, Wnt/β-catenin signaling, TGF-β/Treg differentiation, stemness, and cellular senescence. In AhR pathway, *AHR* expression was significantly enriched in CD4⁺ T-cells from PCS-high individuals; within the TGF-β and Treg signature, a distinct upregulation of multiple genes associated with regulatory T cell function was observed in PCS-high samples, including *JUNB*, *KLF10*, *SMURF2*, *FURIN*, consistent with transcriptional bias toward Treg polarization. In the Wnt/β-catenin signaling module, PCS-high CD4⁺ T-cells showed elevated expression of *SATB1*, *FZD6*, *GSK3B*, *WNT1*, *CTBP2*, and *DVL1*, suggesting active Wnt pathway engagement. The T cell exhaustion signature was also prominently enriched in high PCS donors, including elevated expression of inhibitory receptors *PDCD1* (*PD-1*), *TOX*, *TIGIT*, and *LAG3*. Consistent with a quiescent or less differentiated state, the stemness signature was enhanced in PCS-high samples with increased expression of *BACH2*, *FOXO1*, *FOXO3*, and *TCF7*, key transcription factors that regulate T cell memory and self-renewal. Finally, the senescence signature revealed strong enrichment of *JUN*, *GPX4*, *GSR*, *CDKN1A* (*p21*), and *LMNB1* in PCS-high CD4⁺ T-cells supporting the hypothesis that PCS drives a senescence-like transcriptional program ex vivo. Together, these data validate our prior flow cytometry findings and demonstrate that cell-associated PCS levels stratify CD4⁺ T-cells into distinct transcriptional states. High PCS levels are associated with regulatory and senescence-like profiles, while low PCS levels correlate with effector and cytotoxic phenotypes. Together, these data support a model in which cell-associated PCS influences metabolic remodeling alongside phenotypic and gene expression programs in CD4⁺ T cells ex vivo.

**Figure 4.**
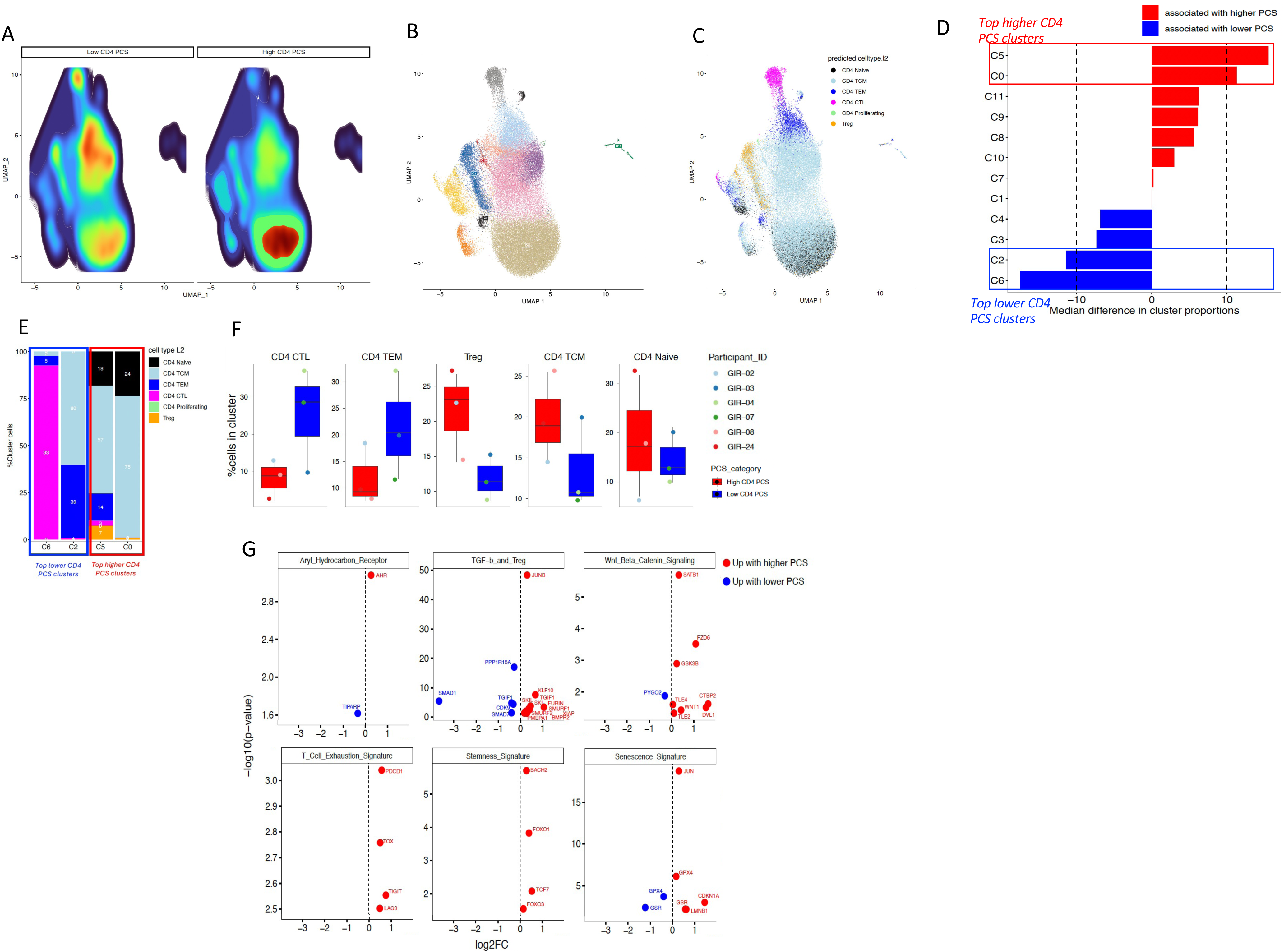
Single-cell transcriptomic analysis of CD4⁺ T-cells from participants with high and low cell-associated PCS concentrations. (**A**) UMAP density plots of CD4⁺ T-cells from six individuals with low (left) or high (right) cell-associated PCS levels. (**B**) UMAP projection of all CD4⁺ T-cells colored by unsupervised cluster assignment. (**C**) UMAP projection colored by predicted CD4⁺ T-cell subset identity. (**D**) Median difference in cluster proportions between high PCS and low PCS groups. (**E**) Stacked bar plot of predicted CD4⁺ T-cell subset identity in clusters C2, C6, C5, and C0, grouped by PCS category. (**F**) Percentage of CD4⁺ T-cell subsets within clusters C2, C6, C5, and C0 shown by individual donor. (**G**) Volcano plots showing differentially expressed genes (red: higher in high PCS; blue: higher in low PCS) within indicated gene modules.

### PCS modulates the transcriptomic and the proteomic prolife of CD4^+^ T-cells in vitro

To determine the functional and molecular impact of PCS on CD4⁺ T-cells as well as the mechanism of action, we conducted series of in vitro assays. Peripheral blood mononuclear cells (PBMCs) were stimulated with anti-CD3/CD28 in the presence of increasing concentrations of PCS. As shown (**Figure 5A**), PCS impaired CD4⁺ T-cell proliferation in a dose-dependent manner, as evidenced by a progressive increase in the proportion of non-proliferating (CTV^high^) cells and a corresponding reduction in proliferating (CTV^low^) cells at 100 μM PCS and as previously assessed (8). The experimental workflow is summarized in **Figure 5B**: CD4⁺ T-cells from five healthy donors were treated with 0, 10, 50, or 100 μM PCS, stimulated for 6 days, and sorted into proliferating and non-proliferating subsets for downstream RNA sequencing and proteomic analysis. Cytokine profiling was also performed on cell culture supernatants after 12h, 24h, day 6 of cell culture. Principal component analysis (PCA) of both transcriptomic and proteomic data revealed clear segregation by PCS concentration and proliferation status, indicating that PCS induces coordinated and dose-dependent molecular remodeling at both transcriptional and protein levels (**Figure 5C**). Transcriptomic analysis of CTV^high^ (non-proliferating) and CTV^low^ (proliferating) CD4⁺ T-cells treated with graded concentrations of PCS revealed broadly similar gene expression patterns across both subsets (**Supplementary Figure 3**). The consistent transcriptional response observed in both populations supports the conclusion that PCS exerts comparable effects irrespective of proliferation status. Accordingly, we focused our in-depth transcriptomic and proteomic analyses on the proliferating CD4⁺ T-cell population.

**Figure 5.**
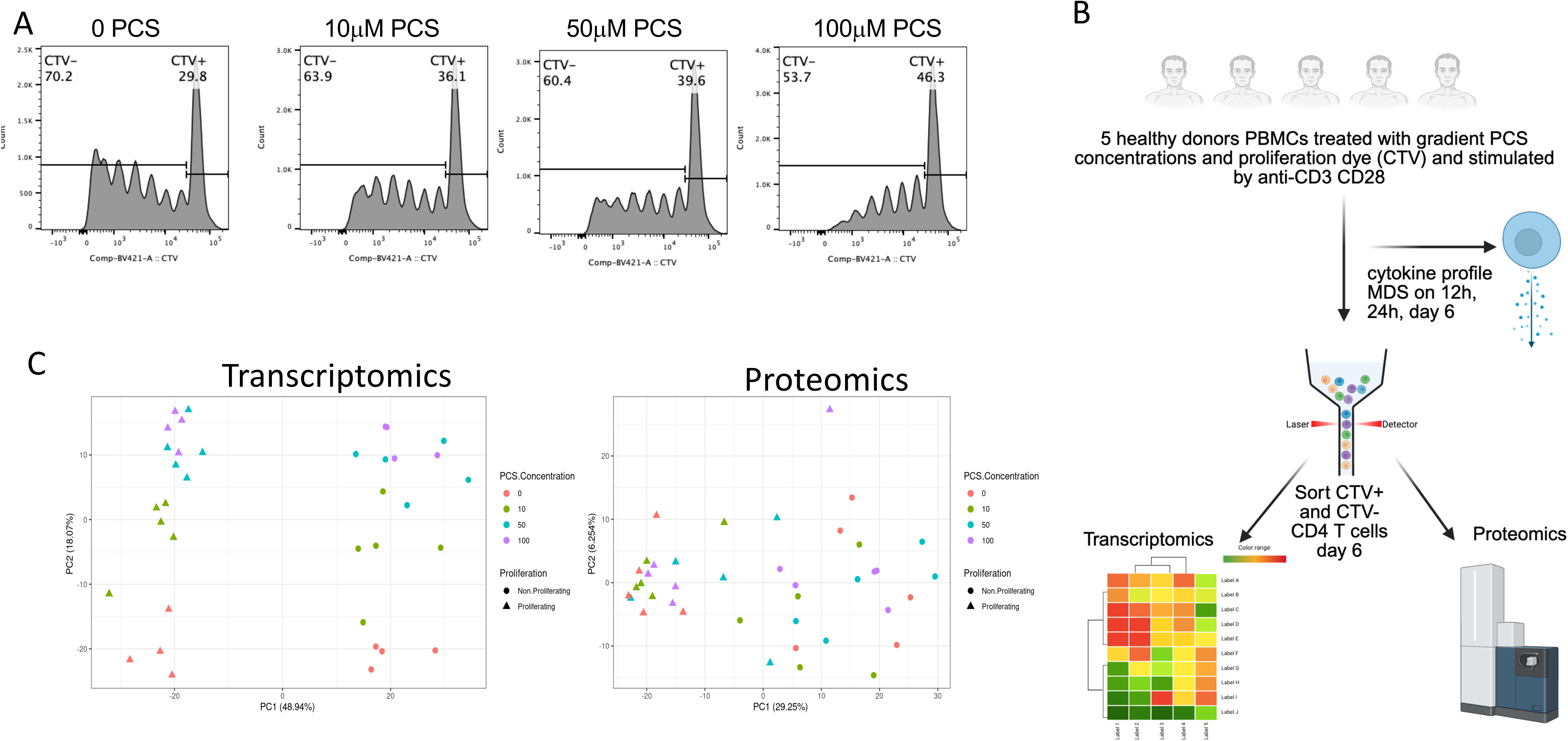
PCS modulates CD4⁺ T-cell proliferation and induces coordinated transcriptomic and proteomic remodeling. **(A)** Representative histograms showing proliferation of CD4⁺ T-cells labeled with CellTrace Violet (CTV) and stimulated with anti-CD3/CD28 in the presence of increasing concentrations of PCS (0, 10, 50, or 100 μM) for 6 days. CTV^high^ (non-proliferating) and CTV^low^ (proliferating) populations are indicated. **(B)** Schematic of the experimental workflow. PBMCs from five healthy donors were labeled with CTV and stimulated with anti-CD3/CD28 in the presence of increasing PCS concentrations. On day 6, CD4^+^ T-cells were sorted for CTV^high^ and CTV^low^ and subjected to transcriptomic and proteomic analysis. Supernatants were collected for cytokine profiling (MDS), and CD4⁺ T-cells were sorted into proliferating and non-proliferating subsets for bulk RNA-seq and proteomic analysis. **(C)** Principal component analysis (PCA) plots of transcriptomic (left) and proteomic (right) datasets from sorted CD4⁺ T-cell populations. Samples cluster by PCS concentration and proliferation status, indicating that PCS induces dose-dependent molecular remodeling across both omics layers.

### PCS Induces a senescence program in CD4⁺ T-cells in vitro

To evaluate the effect of PCS on proliferating CD4⁺ T-cells, we performed bulk RNA sequencing on sorted CTV^low^ CD4⁺ T-cells stimulated in vitro for 6 days with anti-CD3/CD28 in the presence of increasing PCS concentrations (0–100 μM). PCS treatment led to dose-dependent activation of the AhR pathway (*AHR*, *AHRR, CYP1B1, TIPARP*), suggesting that PCS engages xenobiotic response signaling (**Figure 6A**) and confirming the ex vivo data obtained by scRNA-seq (**Figure 4G**). In parallel, TGF-β/Treg signaling pathway (*SMADs, FOXP3*) were also upregulated (**Figure 6B**), consistent with the induction of Treg. PCS exposure also suppressed the expression of key glycolytic genes (*HK1, LDHA, GAPDH*) (**Figure 6C**) and mTOR pathway components (*RPS6, EIF4EBP1, S6K*) (**Figure 6D**), indicating profound metabolic reprogramming and inhibition of biosynthetic activity. PCS induces Notch signaling pathway (**Figure 6E**), and the Wnt/β-catenin signaling pathway (**Figure 6F**), with increased expression of *LEF1, TCF7, FZD* family genes. Genes associated with T cell stemness, including *fOXO1, FOXO3, LEF1,* and *TCF7*, were progressively upregulated with higher PCS concentrations (**Figure 6G**). Meanwhile, PCS induced prominent expression of T cell exhaustion markers such as *PDCD1, TOX,* and *LAG3* (**Figure 6H**), features typically associated with chronic stimulation and loss of effector function. Finally, genes linked to immune senescence, including *CDKN1A (p21), CDKN2A (p16), JUN,* and *CD69*, were markedly upregulated at higher PCS doses (**Figure 6I**), confirming the acquisition of a senescent-like transcriptional profile.

**Figure 6.**
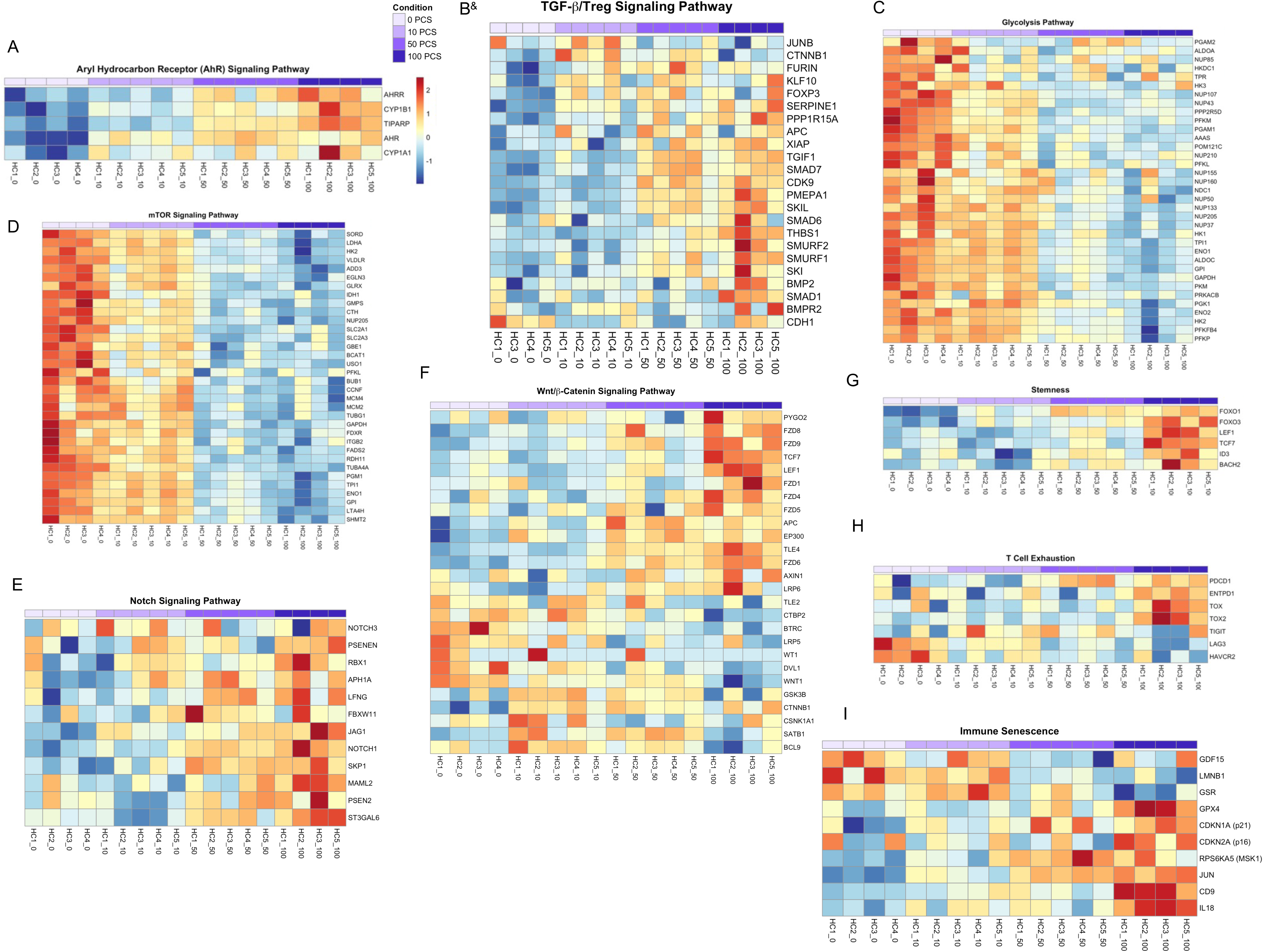
PCS induces transcriptomic reprogramming in proliferating CD4⁺ T-cells. Bulk RNA sequencing was performed on sorted CTV^low^ CD4⁺ T-cells after 6 days of in vitro stimulation with anti-CD3/CD28 in the presence of 0–100 µM PCS. Heatmaps display normalized gene expression (z-scores) across treatment conditions for selected pathways and gene sets. Panel (**A**) shows genes from the AhR signaling pathway, (**B**) TGF-β-Treg signaling pathway, (**C**) glycolysis pathway, (**D**) mTOR signaling pathway, (**E**) Notch signaling pathway, (**F**) Wnt/β-catenin signaling pathway, (**G**) stemness-associated genes, (**H**) T cell exhaustion markers, and (**I**) immune senescence markers. Each column represents one sample, and each row corresponds to an individual gene. Expression levels are color-coded from low (blue) to high (red). ^&^ Indicate that this heatmap was generated from non-proliferating CTV^high^ cells.

Together, the transcriptional programs induced by PCS in vitro by bulk RNA-seq closely mirror the ex vivo scRNA-seq signatures from high-PCS donors, indicating that PCS drives a consistent CD4⁺ T-cell state across experimental systems. The in vitro experiments conducted to elucidate the mechanism of action of PCS revealed activation of similar transcriptional and phenotypic programs, such as senescence, Treg differentiation, Wnt/β-catenin signaling, as those identified by scRNA-seq analysis of ex vivo CD4⁺ T-cells from PCS-high participants. These transcriptional changes reveal that PCS orchestrates a coordinated program in CD4⁺ T-cells characterized by xenobiotic sensing, metabolic remodeling and inhibition, stem-like adaptation, immune exhaustion, and convergence toward a senescence-like state.

To investigate the phenotypic impact of PCS exposure during CD4⁺ T-cell activation and to validate the transcriptomic signature, we cultured human CD4⁺ T-cells in vitro with anti-CD3/CD28 in the presence of increasing concentrations of PCS (0, 50, 100 µM). Flow cytometry analysis of CCR7 vs. CD45RA (top panels, **Figure 7A**) showed a progressive redistribution of T cell subsets. Specifically, the frequency of CCR7⁺CD45RA⁻ TCM cells increased from 27.4.3 ± 3.6% at 0 µM PCS to 37.0 ± 2.6% at 50 µM and 42.0 ± 3.16% at 100 µM. Meanwhile, TEMRA (CCR7⁻CD45RA⁺) cells decreased from 4.8 ± 1.2% (0 µM) to 3.1 ± 0.5% (50 µM) and 1.2 ± 0.4% (100 µM), consistent with the PCS-associated TEMRA contraction observed ex vivo (**Figure 3D**). TEM cells declined (30.9 ± 13.8%, 26.4 ± 14.4%, 21.4 ± 11.6%), while naïve cell frequencies remained relatively stable.

**Figure 7.**
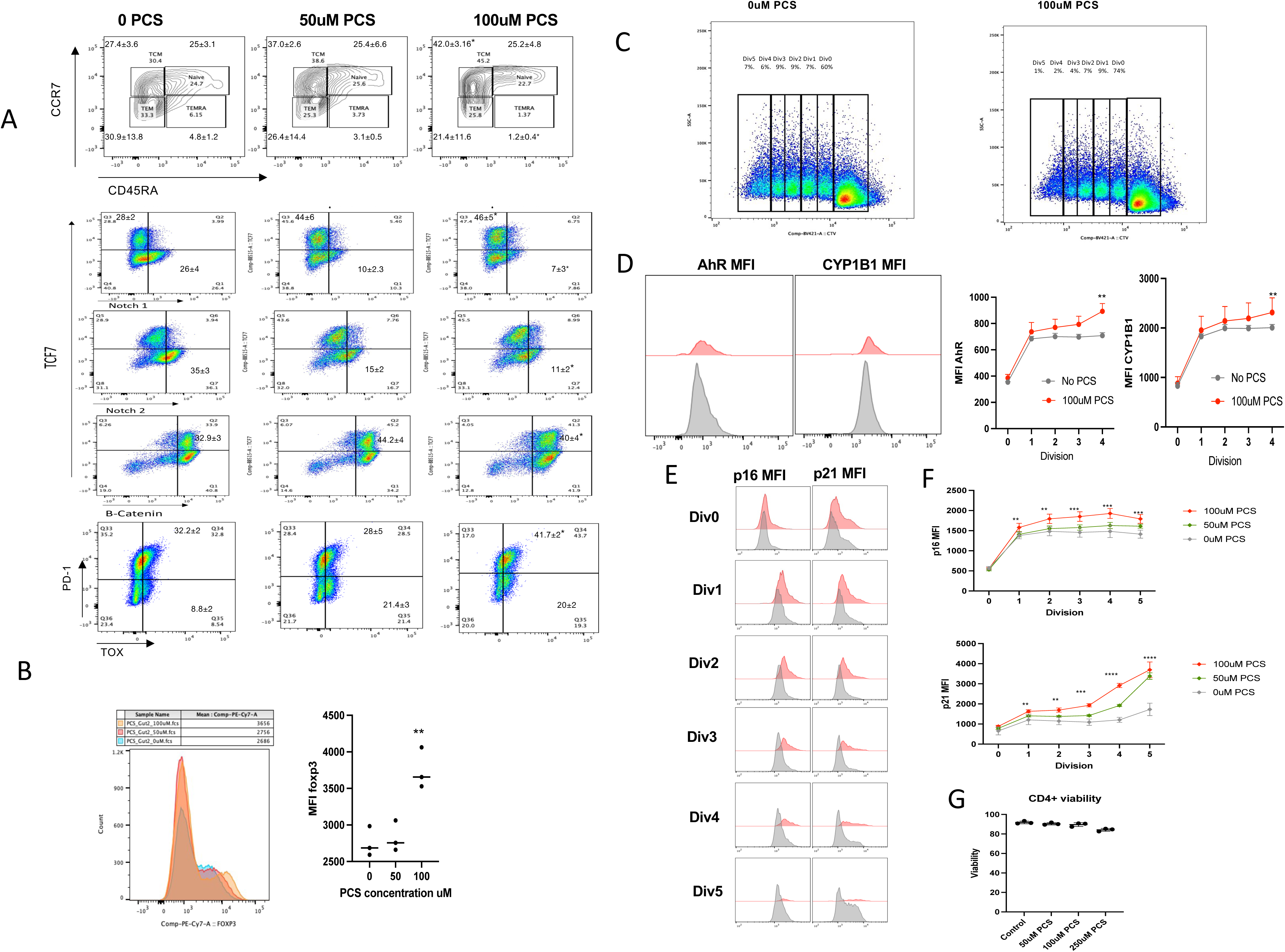
PCS induces senescence markers and alters CD4⁺ T-cell phenotype in vitro. **(A**) Representative flow cytometry plots of CD4⁺ T-cells stimulated with anti-CD3/CD28 for 6 days in the presence of 0, 50, or 100 μM PCS, showing CCR7 vs. CD45RA (top panels) and TCF7, Notch1, Notch2, β-catenin, and PD-1 vs. TOX (middle and bottom panels). Values in quadrants indicate median ± SD across 3–5 biological replicates. (**B**) Overlay histogram (left) and quantification (right) of FOXP3 expression in naïve CD4⁺ T-cells cultured in the presence of IL-2 and increasing concentrations of PCS (0, 50, or 100 μM) for 5 days. (**C**) CellTrace Violet dilution plots showing proliferative divisions of CD4⁺ T-cells cultured with or without 100 μM PCS. (**D**) MFI Flow cytometry histograms (left) and quantification (right) of AhR and CYP1B1 expression across divisions (Div0–Div4) for each PCS condition. (**E**) Flow cytometry histograms of p16 and p21 expression across divisions (Div0–Div5) for each PCS concentration**. (F**) Quantification of p16 and p21 MFI across cell divisions in each PCS condition. (**G**) Viability of CD4⁺ T-cells after 6 days of culture with or without PCS, assessed by fixable viability dye. Statistical comparisons were performed using one-way ANOVA with multiple comparisons correction: *p < 0.05; **p < 0.01; ***p < 0.001; ****p < 0.0001.

In the middle and lower panels, PCS promoted a strong dose-dependent induction of TCF7 (28 ± 2% → 44 ± 6% → 46 ± 5%) and β-catenin (32.9 ± 3% → 44.2 ± 4% → 40.0± 4%) expression, supporting the activation of the Wnt signaling pathway. Additionally, Notch1 and Notch2 surface expression was reduced (Notch1⁺: 26 ± 4% → 10 ± 2.3% → 7 ± 3%; Notch2⁺: 35.3 ± 3% → 15 ± 2% → 11 ± 2%), indicating ligand engagement and downstream signaling. These phenotypic changes parallel the transcriptional activation of the Wnt/β-catenin and Notch pathways identified in RNA-seq analysis (**Figure 6D–F**), and altogether reflect a PCS-driven shift toward a regulatory or non-effector CD4⁺ T-cell state.

In vitro assays demonstrated that PCS directly upregulates FOXP3 protein expression (**Figure 7B**). Stimulation of naïve CD4⁺ T-cells with anti-CD3/CD28 in the presence of PCS led to a clear dose-dependent induction of FOXP3⁺ cells. These data validate the Treg induction detected in the ex vivo flow cytometric (**Fig. 3C, 3D)** and in the scRNA-seq (**Fig. 4F, and 4G**) analyses.

To validate AhR pathway activation suggested by the transcriptomic analysis (**Fig. 4G**, and **Fig 6A**), we next assessed AhR protein expression and its canonical downstream target CYP1B1 in CD4⁺ T-cells cultured with or without PCS. Flow cytometry analysis across proliferative divisions (Div0–Div4) **(Figure 7C)** revealed a progressive increase in both AhR and CYP1B1 median fluorescence intensity (MFI) with increasing PCS concentrations (**Figure 7D**). Cells exposed to 100 µM PCS displayed a significant elevation in AhR and CYP1B1 expression beginning at the first division and persisting through later divisions, compared to untreated controls (p < 0.01). These findings confirm that PCS activates the AhR signaling pathway in proliferating CD4⁺ T-cells, consistent with the observed upregulation of AhR-related genes (including CYP1B1 and AhR) in transcriptomic datasets.

To assess senescence-related responses, we quantified p16 and p21 expression across cell divisions, tracked by CTV dilution (**Figures 7C and 7E**). PCS markedly increased p16 and p21 expression in proliferating CD4⁺ T-cells, with the highest MFI detected in late divisions. Quantitative analysis confirmed a significant, dose-dependent elevation of both markers in PCS-treated cells across all divisions (**Figures 7E–7F**). Importantly, PCS treatment did not compromise cell viability (**Figure 7G**), indicating that reduced proliferation was not attributable to cytotoxicity.

The aggregate findings show that PCS reprograms CD4⁺ T-cell differentiation, metabolic remodeling, and signaling by activating Wnt/β-catenin and Notch pathways, enhancing FOXP3 expression, and inducing senescence-associated proteins. These changes integrate with transcriptomic evidence (**Figure 4** and **Figure 6**) to support a model in which PCS imposes a non-proliferative, regulatory-like, and senescent transcriptional state that may contribute to long-term T cell dysfunction and immune cell aging.

### PCS Induces a Proteomic Shift from Immune Effector Programs to Mitochondrial and Stress-Adapted States in CD4⁺ T-Cells in vitro

To define the impact of PCS on CD4⁺ T-cell function at the protein level, we examined global proteomic changes across a PCS concentration gradient (0–100 μM). We identified a coordinated pattern of downregulation of immune-associated proteins and upregulation of mitochondrial, metabolic, and stress-related proteins, suggesting broad functional remodeling under PCS exposure (**Figure 8**). A set of proteins involved in T cell signaling, cytotoxic function, and intracellular trafficking were significantly downregulated with increasing PCS concentration: FCHO1(47), an adaptor protein critical for clathrin-mediated endocytosis, particularly involved in TCR internalization and recycling, essential for antigen sensitivity; SLC39A8 (ZIP8), a zinc transporter responsible for maintaining zinc homeostasis, which is necessary for T cell activation, cytokine signaling, and redox defense(48); SORL1, a receptor involved in endosomal trafficking and membrane protein recycling(49), potentially affecting surface receptor turnover and antigen presentation; OTUD6B, a deubiquitinase that regulates protein stability and immune signaling pathways(50), potentially influencing T cell fate decisions and inflammation control; ADAM8, a metalloproteinase involved in leukocyte adhesion, migration, and tissue infiltration(51); its downregulation may impair immune surveillance; GZMA (Granzyme A), a cytototoxic effector molecule secreted by activated T cells to induce target cell apoptosis(52); MAPK13 (p38δ), member of the p38 MAPK family, key for inflammatory signaling, stress responses, and T cell polarization(53). These results indicate that PCS suppresses proteins central to immune effector activity, antigen receptor signaling, and genomic stress protection, suggesting compromised immune function under PCS burden.

**Figure 8.**
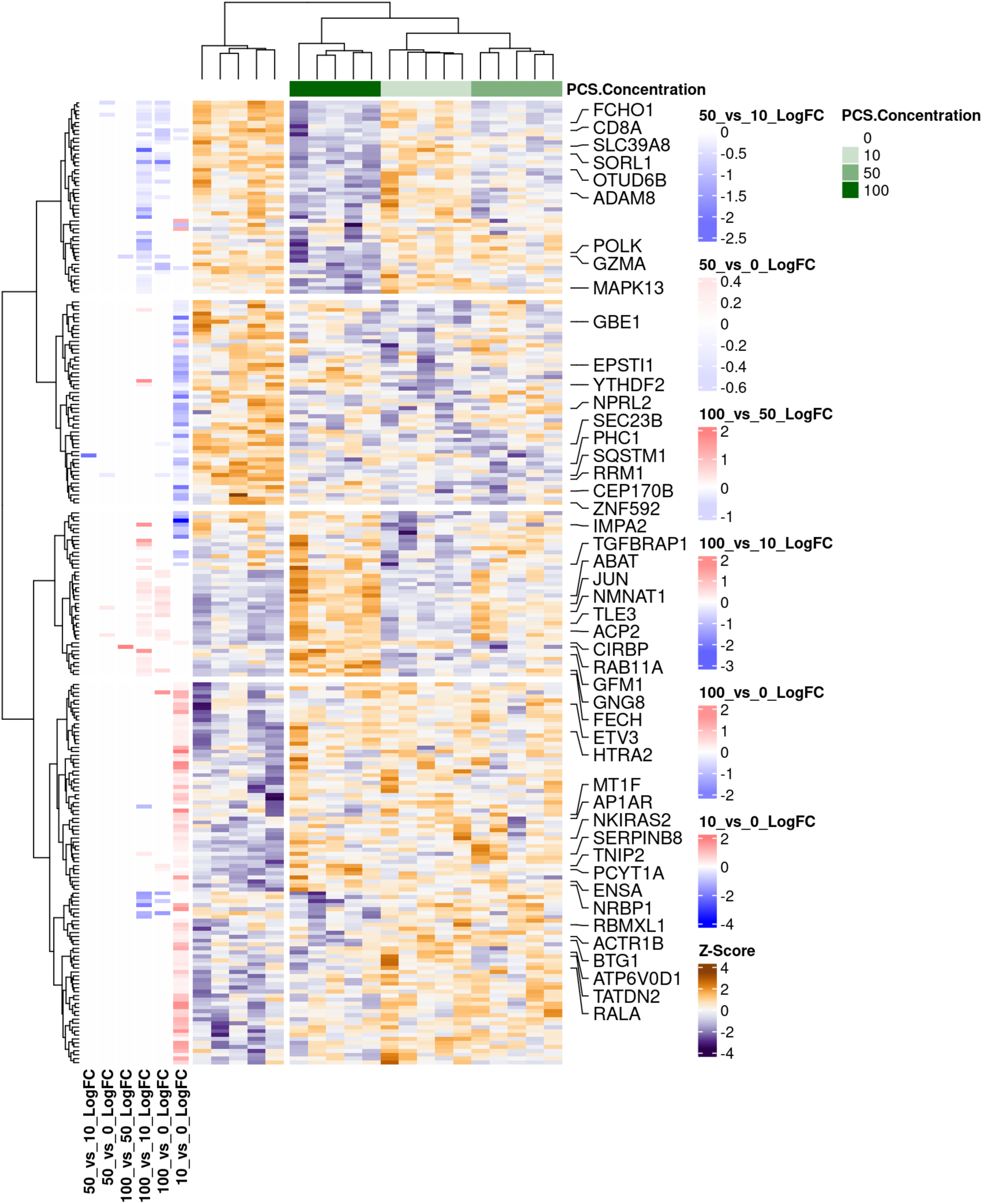
Proteomic profiling of proliferating CD4⁺ T-cells reveals dose-dependent remodeling induced by PCS. Heatmap depicting z-score normalized expression of differentially abundant proteins in CTV^low^ proliferating CD4⁺ T-cells exposed to a gradient PCS concentration (0, 10, 50, and 100 μM). Unsupervised hierarchical clustering highlights protein expression patterns across conditions. Proteins involved in T cell activation and cytotoxicity (e.g., GZMA, MAPK13) and signaling regulation (e.g., SORL1, FCHO1, OTUD6B) are progressively downregulated with increasing PCS exposure. In contrast, proteins linked to metabolic stress adaptation, mitochondrial function, and transcriptional regulation (e.g., NMNAT1, TGFBRAP1, ABAT, JUN, GFM1, CIRBP) are upregulated. Side heatmaps show log₂ fold changes for pairwise comparisons between PCS conditions.

In contrast, several proteins were upregulated in response to higher PCS levels, forming distinct clusters associated with metabolic adaptation, transcriptional regulation, and mitochondrial stress responses: 1) Mitochondrial Function & Oxidative Stress Adaptation, GFM1, a mitochondrial translation factor, indicating enhanced mitochondrial protein synthesis(54), possibly compensating for stress; FECH, an enzyme catalyzing the final step of heme biosynthesis, often upregulated in response to mitochondrial dysfunction(55); HTRA2, a mitochondrial serine protease involved in mitochondrial quality control and apoptosis signaling(56), RAB11A, a GTPase regulating endosomal recycling, potentially involved in mitochondrial and vesicle trafficking(57); GNG8, a G-protein subunit involved in signal transduction(58), possibly compensating for upstream signaling defects. 2) Transcriptional and Post-Transcriptional Regulation: JUN, a transcription factor within the AP-1 complex, upregulated in response to cellular stress(59); ETV3, transcriptional repressor that modulates immune activation and inflammation resolution(60); TLE3, a corepressor that interacts with transcription factors in Wnt and Notch signaling, potentially regulating T cell differentiation(61); CIRBP, an RNA-binding protein involved in mRNA stability and translation under stress conditions(62); TGFBRAP1, a TGF-β signaling adaptor linking to SMAD pathways(63), suggesting activation of immunoregulatory responses. 3) Metabolic Remodeling and Stress Buffering proteins: NMNAT1, an enzyme critical for NAD⁺ biosynthesis, supporting redox balance and energy metabolism(64); ACP2, a lysosomal acid phosphatase involved in catabolic processes and cellular stress adaptation(65); ABAT, a GABA transaminase, potentially linked to mitochondrial metabolism and cellular quiescence(66); IMPA2, involved in inositol phosphate metabolism, important for second messenger signaling and metabolic regulation(67). These proteins collectively reflect a compensatory shift toward a metabolically stress-adapted state, potentially at the expense of immune function and transcriptional responsiveness.

Altogether, the proteomic profiling suggests that PCS leads to the downregulation of key immune effector and signaling proteins, including those responsible for TCR signaling, zinc transport, cytotoxicity, and inflammation. In parallel, PCS promotes the upregulation of mitochondrial, metabolic, and stress-adaptive proteins, indicative of a transcriptionally reprogrammed, energetically burdened, and functionally suppressed T cell state.

### PCS Suppresses Cytokine Production and Impairs Th1/Th2 Polarization in CD4⁺ T-Cells

To evaluate the effect of PCS on cytokine production and T helper cell polarization, we profiled 40 cytokines in supernatants from PBMCs stimulated with anti-CD3/CD28 in the presence of increasing concentrations of PCS (0, 10, 50, and 100 μM) over 12 hours, 24 hours, and 6 days. Heatmap analysis revealed dose- and time-dependent modulation of inflammatory, homeostatic, and regulatory cytokines across all five donors (**Supplementary Figure 4**). Unsupervised clustering identified several cytokine modules selectively affected by PCS exposure (**Supplementary Figure 4B**). Notably, Cluster 1, including *IL-7*, *SDF-1a*, and *TGF-β3*, was transiently upregulated at 24 hours with 100 μM PCS (*P* = 0.042), whereas Clusters 2, 4, 6, and 7, encompassing pro-inflammatory (*IL-6*, *TNF-α*) and Th-associated (*IL-4*, *IL-5*, *IL-13*, *IL-17A*) cytokines, were consistently downregulated across multiple time points. These findings suggest that PCS suppresses both early and late inflammatory cytokine production, including those associated with Th1 and Th2 lineage specification. To assess the impact of PCS on CD4⁺ T-cell lineage differentiation, PBMCs were stimulated under non-polarizing, Th1-polarizing, or Th2-polarizing conditions in the presence or absence of 100 μM PCS for 6 days. Flow cytometric analysis was performed on gated CD4⁺ T-cells to evaluate the expression of GATA3 (Th2 master regulator) and T-bet (Th1 master regulator). Representative contour plots and mean fluorescence intensity (MFI) values are shown (**Supplementary Figure 4C**). In non-polarized conditions, PCS treatment led to a marked reduction in GATA3 expression (MFI decreased from 2370 to 2018) and T-bet expression MFI decreased from 4166 to 2018), indicating that PCS suppresses basal expression of lineage-specifying transcription factors in CD4⁺ T-cells. Under Th2-polarizing conditions, PCS exposure significantly impaired Th2 commitment, as demonstrated by a substantial reduction in both the frequency and MFI of GATA3⁺ CD4⁺ T- cells. GATA3 MFI dropped from 6688 (no PCS) to 2003 (with PCS), accompanied by a decrease in the CD45RA⁻ GATA3⁺ population from 40.0% to 28.7%, suggesting that PCS limits effective Th2 polarization. Similarly, in Th1-polarizing conditions, PCS reduced the frequency and expression level of T-bet⁺ CD4⁺ T-cells. T-bet MFI decreased from 3754 to 2576, with a corresponding reduction in the T-bet⁺ CD45RA⁻ subset from 37.3% to 37.0%. While the frequency change was modest, the decrease in MFI suggests blunted transcriptional programming of Th1 cells under PCS exposure.

Altogether, our comprehensive multi-omics analysis reveals that PCS exerts profound and coordinated effects on CD4⁺ T-cell biology. Ex vivo and in vitro transcriptomic profiling of CD4⁺ T-cells exposed to PCS uncovered a shift toward immunoregulatory and stem-like states, with activation of TGF-β, Wnt/β-catenin, and AhR signaling pathways, alongside suppression of mTOR, and glycolysis. These transcriptional changes were mirrored at the proteomic level by downregulation of proteins essential for T cell signaling, cytotoxicity, and antigen trafficking, and by upregulation of proteins involved in mitochondrial maintenance, metabolic stress adaptation, and transcriptional repression. Functionally, PCS impaired CD4⁺ T-cell proliferation, suppressed pro-inflammatory and lineage-defining cytokines, and blunted Th1 and Th2 polarization, as shown by reduced expression of the transcription factors T-bet and GATA3. Together, these findings suggest that PCS reprograms CD4⁺ T-cells into a metabolically restrained and functionally suppressed state, characterized by induction of senescence program that leads to CD4^+^ T-cell immune cell aging characteristics of immune cells in PLWH.

## Discussion

In this study, we show that GDBMs remodel metabolic pathways in human CD4⁺ T-cells and reshape key aspects of their cellular biology. Integrating ex vivo metabolomics with scRNA-seq, bulk RNA-seq, and high-dimensional cytometry (t-SNE/FlowSOM), we identify a coherent program of metabolic remodeling, senescence induction, and diminished effector potential driven by cell-associated PCS, positioning this GDBM as a key cell-intrinsic regulator of CD4⁺ T-cell aging. Our findings culminate in a model in which the GDBM PCS acts as a central driver of CD4⁺ T-cell senescence. As illustrated in **Figure 9**, our integrated analyses support a mechanistic model in which PCS engages an AhR-associated signaling axis that coordinates metabolic remodeling with transcriptional reprogramming in CD4⁺ T-cells. This PCS–AhR axis is associated with induction of TGF-β signaling alongside suppression of glycolytic and mTOR-dependent anabolic pathways, key features of metabolic remodeling and anabolic arrest that constrain effector T cell proliferation and function. In parallel, PCS exposure coincides with enhanced TGF-β pathway activity, increased FOXP3 expression, and acquisition of regulatory-like features. Concurrent activation of Wnt/β-catenin signaling and marked upregulation of TCF7 further point toward a stem-like, immune-adapted cellular state. Importantly, PCS is also associated with robust induction of the cell-cycle inhibitors p16 and p21, consistent with proliferative arrest and a shift toward cellular senescence. Together, these metabolic and transcriptional alterations converge on a phenotype characterized by metabolic quiescence, transcriptional restraint, and impaired proliferative capacity, promoting CD4⁺ T cell immune aging characteristic of PLWH. Given that AhR is a cytoplasmic ligand-activated transcription factor(68), the induction of AhR-responsive programs by PCS is consistent with intracellular engagement of PCS.

**Figure 9.**
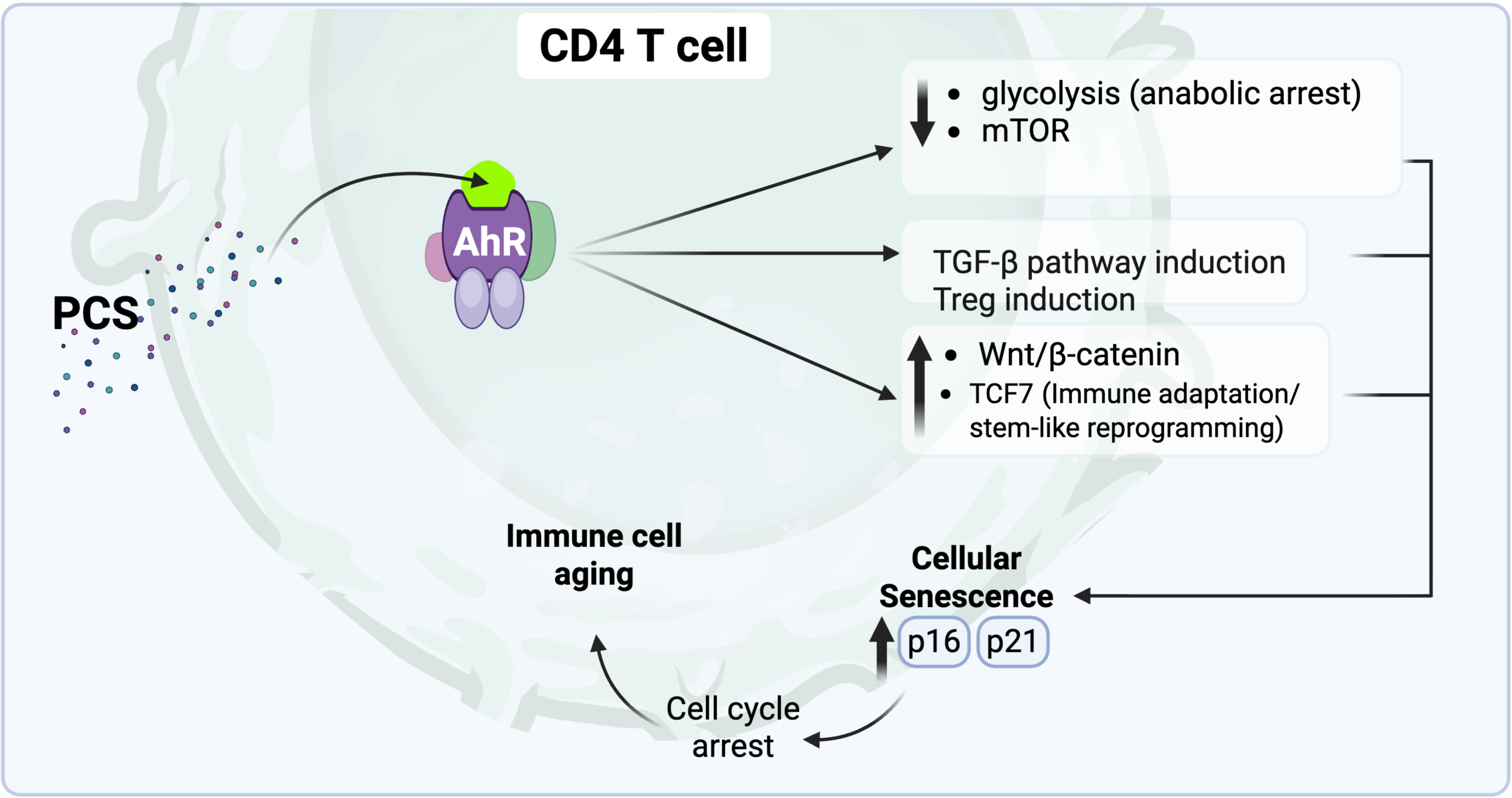
Cell-associated PCS engages AhR-associated signaling pathways in CD4⁺ T cells, coordinating metabolic and transcriptional reprogramming. PCS-associated activation of AhR-responsive programs coincides with suppression of glycolysis and mTOR signaling, consistent with anabolic arrest. In parallel, PCS exposure is associated with enhanced TGF-β signaling and increased FOXP3 expression, indicative of regulatory-like differentiation. Concurrent activation of Wnt/β-catenin signaling and upregulation of TCF7 further point toward a stem-like, immune-adapted cellular state. These coordinated signaling and metabolic changes are accompanied by induction of the cell-cycle inhibitors p16 and p21, consistent with proliferative arrest and cellular senescence. Together, these processes converge on a non-proliferative, metabolically restrained state that promotes CD4⁺ T-cell immune aging characteristic of PLWH.

Our observation that PCS induces expression of canonical senescence markers p16 and p21 CD4⁺ T-cells aligns with growing evidence that immune cells, particularly T cells, undergo senescence in both aging and chronic disease. Previous studies have identified p16 and p21 as hallmarks of aged T-cells ex vivo, associated with reduced proliferative capacity, altered metabolic programming, and increased susceptibility to functional exhaustion (21, 23, 24). These findings suggest that PCS likely acts as an upstream inducer of immunosenescence, establishing a transcriptional and metabolic program reminiscent of aged immune cells. Thus, PCS exposure may not only reflect, but also potentially exacerbate, aging-associated immune decline characteristic of PLWH (25, 31, 69).

Emerging evidence suggests that AhR may serve as a central molecular link between environmental/metabolic cues and immune-cell aging. AhR is broadly expressed in immune cells and integrates signals from dietary, microbial, and endogenous ligands to regulate gene expression programs relevant to cell differentiation, metabolism, and immune function (68, 70, 71). Mechanistically, AhR influences mitochondrial function, proteostasis, and reactive oxygen species balance, processes intimately connected to the hallmarks of cellular aging (71, 72). In the context of immune cells, recent reviews have proposed that AhR’s regulation of differentiation and metabolic programs may predispose lymphocytes to aging-like phenotypes when chronically engaged by endogenous or microbial metabolites (70), (73). Given our observation that PCS (a GDBM) robustly induces AhR and downstream signaling in CD4⁺ T-cells, and along with induction of senescence markers (p16, p21), mitochondrial and metabolic dysfunction, our data provide a plausible mechanistic route by which chronic AhR engagement drives immunosenescence. This positions AhR not just as a xenobiotic sensor, but also as a modulator of immune cell homeostasis and longevity, a concept that may have broad implications for chronic infection, aging, and immune-mediated diseases.

Long-lived and transcriptionally inert CD4⁺ T-cell subsets are recognized as critical components of the latent HIV-1 reservoir that persists despite prolonged ART. Among these, T memory stem cells (TSCM) represent a uniquely durable viral reservoir. Buzon et al. demonstrated that TSCM cells harbor the highest per-cell levels of HIV-1 DNA compared to other memory subsets, retain replication-competent virus, and exhibit remarkable reservoir stability over nearly a decade of suppressive ART (74). These findings highlight the ability of HIV-1 to exploit the stem-like qualities of TSCM self-renewal, resistance to apoptosis, and multipotency, to evade eradication. Extending this concept, a study (75) identified senescence-like transcriptional programs in central memory CD4⁺ T-cells of ART-treated individuals with poor immune recovery. These senescent cells displayed elevated TGF-β signaling, and oxidative stress responses traits that were positively correlated with inducible HIV reservoirs. Foundational studies by Finzi et al.(76) and Chomont et al (45) further established that HIV preferentially resides in long-lived memory T cells. Together, these findings suggest a unifying model in which HIV-1 capitalizes on both stemness and senescence within the CD4⁺ T-cell compartment phenotypes that confer resistance to cytolytic mechanisms, immune surveillance, and apoptosis. Our data corroborate these findings demonstrating that PCS induces CD4⁺ T-cell senescence characterized by increased expression of p16, p21, and exhaustion markers, a cell state permissive for HIV-1 reservoir maintenance.

The accumulation of metabolic intermediates is often linked to dysfunctional metabolic pathways. Under normal conditions, metabolic intermediates are efficiently processed through enzymatic reactions to sustain cellular homeostasis. When a pathway is disrupted due to enzyme dysfunction, altered signaling, oxidative stress, or mitochondrial defects, intermediates may build up, leading to metabolic stress and cellular dysfunction. As examples, succinate accumulation in macrophages leads to HIF-1α stabilization and a pro-inflammatory state (20, 77). Lactate build-up due to impaired oxidative phosphorylation leads to metabolic acidosis (78).TCA cycle intermediates (e.g., citrate, fumarate) accumulating due to mitochondrial dysfunction can alter redox balance and epigenetic regulation (79). The ex vivo correlation network revealed that multiple GDBMs converge on shared metabolic pathways within CD4⁺ T-cells, defining a common immunometabolic signature associated with immune dysfunction (**Figure 1** and **Figure 2**). PAG and PCG have not previously been shown to modulate immune cells; here we detect both metabolites at the cellular level in human CD4⁺ T-cells. PAG exhibited significant positive correlations with multiple metabolic pathways (**Figure 1D**) and with pathways that were themselves positively associated with the size of the intact HIV-1 reservoir (**Figure 2D**). A similar pattern was observed for PCG suggesting its role in modulating CD4⁺ T-cells biology as well. Further studies are needed to address the mechanism of action of PAG and PCG on CD4⁺ T-cells. Our data provide evidence that intact and total HIV-1 reservoirs only partially overlap in their metabolic associations, suggesting distinct metabolic dependencies for HIV-1 reservoir maintenance. As illustrated in **Figure 2D**, only a limited set of metabolites, uridine 5’-diphosphate (UDP), deoxyuridine triphosphate, and bilirubin, were uniquely associated with the total reservoir. In contrast, the intact reservoir displayed broader associations with dysfunctional central carbon and nucleotide metabolism, indicating that metabolite-driven immune cell dysfunction may preferentially support the persistence of intact, replication-competent proviruses. Our findings suggest that the metabolic requirements for HIV-1 reservoir maintenance differ when comparing total versus intact proviruses, with intact HIV-1 reservoirs exhibiting stronger associations with broad immunometabolic dysfunction, while total reservoir size appears linked to a narrower set of metabolic pathways. To our knowledge, this is the first study to perform CD4⁺ T-cell metabolic profiling and show that intact and total HIV-1 proviruses are associated with distinct metabolic signatures, indicating that these two reservoirs may be shaped and maintained through different immunometabolic states.

While our findings demonstrate that cell-associated PCS is linked to extensive metabolic, transcriptional, and phenotypic remodeling of CD4⁺ T cells, several considerations are important for contextualizing these observations. Because our analyses are based on cross-sectional ex vivo profiling, they do not establish the temporal sequence or causality of PCS exposure and CD4⁺ T-cell reprogramming in vivo. In addition, cell-associated PCS levels likely reflect a dynamic balance of cellular uptake, binding, and clearance that was not longitudinally assessed, potentially underestimating transient or fluctuating intracellular exposures. Although our in vitro activation system enabled mechanistic interrogation of PCS-associated pathways, it does not fully capture the complexity of tissue microenvironments, antigenic histories, or stromal and microbial cues that shape CD4⁺ T-cell fate in vivo. Moreover, while PCS served as a mechanistic prototype, other aromatic gut-derived bacterial metabolites that co-associate with similar metabolic pathways, including PAG and PCG, were not functionally interrogated, leaving open the possibility of cooperative or additive effects within this metabolite network. More broadly, these metabolites remain insufficiently explored in immunity, and future studies will be required to define how their cell-associated accumulation influences diverse immune compartments across disease contexts. Finally, interpretation of the observed associations with HIV reservoir size is constrained by the absence of a tractable in vitro system that faithfully models reservoir establishment and long-term persistence. Existing latency and latency-reversal assays primarily assess proviral inducibility and do not recapitulate the metabolic quiescence, cellular longevity, and survival programs that govern reservoir maintenance in vivo. Accordingly, our study emphasizes ex vivo human associations linking cell-associated GDBMs to immunometabolic and senescence-associated CD4⁺ T-cell states previously implicated in HIV persistence, rather than attempting to model reservoir dynamics in vitro. Future studies leveraging improved experimental systems and interventional approaches will be necessary to directly test the causal contribution of microbiome-derived metabolites to reservoir stability.

Our findings support a systems-level model in which GDBMs act as endocrine-like regulators of immune metabolism, influencing the balance between immune activation, quiescence, and senescence. Within this framework, PCS emerges as a prototype effector that recapitulates the broader metabolic dysfunction observed ex vivo, providing a mechanistic entry point to dissect how chronic microbial exposure drives immune aging and shape T-cell homeostasis. Thus, strategies aimed at modulating GDBMs levels, via gut microbiome targeting or dietary interventions, may restore metabolic balance, reverse immune senescence-associated aging, and may disrupt HIV reservoir stability.

---

**Supplementary Figure 1.**
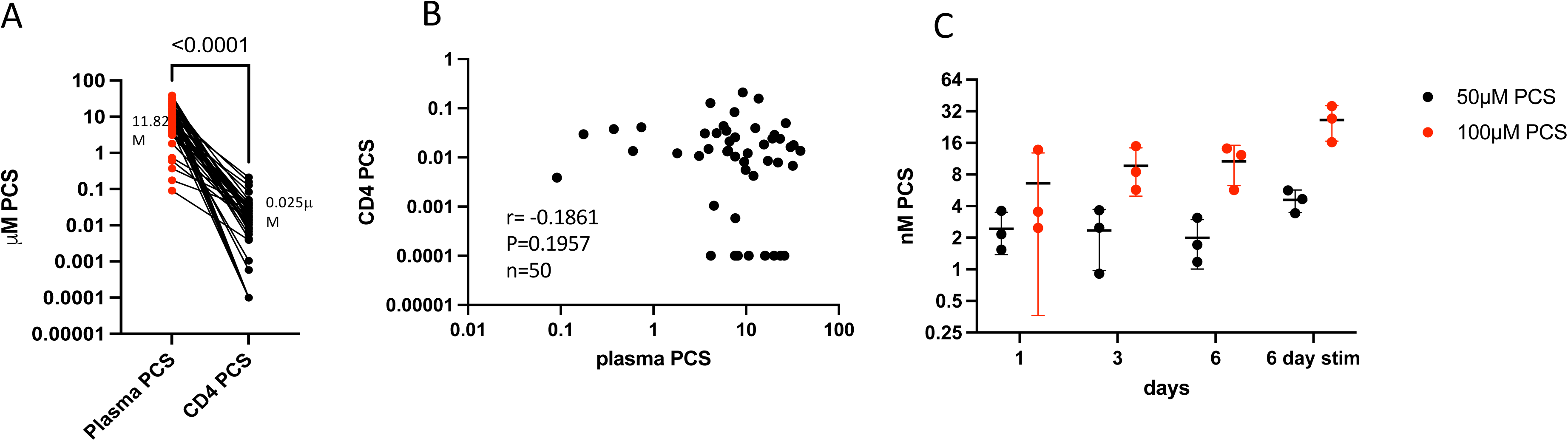
PCS concentrations in plasma and CD4⁺ T-cells from PLWH reveal tissue-specific accumulation and lack of correlation. (**A**) Mass spectrometry was used to quantify PCS concentrations in paired plasma and CD4⁺ T-cell samples from 50 PLWH. The mean PCS concentration in plasma was 11.82 μM, whereas the mean cell-associated concentration in CD4⁺ T-cells was 0.025 μM. (**B**) Spearman correlation analysis between plasma and CD4⁺ T-cell PCS concentrations. (**C**) Cell-associated PCS accumulation in CD4⁺ T-cells is shown from a separate experiment in which cells were incubated in vitro with 50 or 100 μM PCS for 1, 3, or 6 days, with or without TCR stimulation.

**Supplemental Figure 2.**
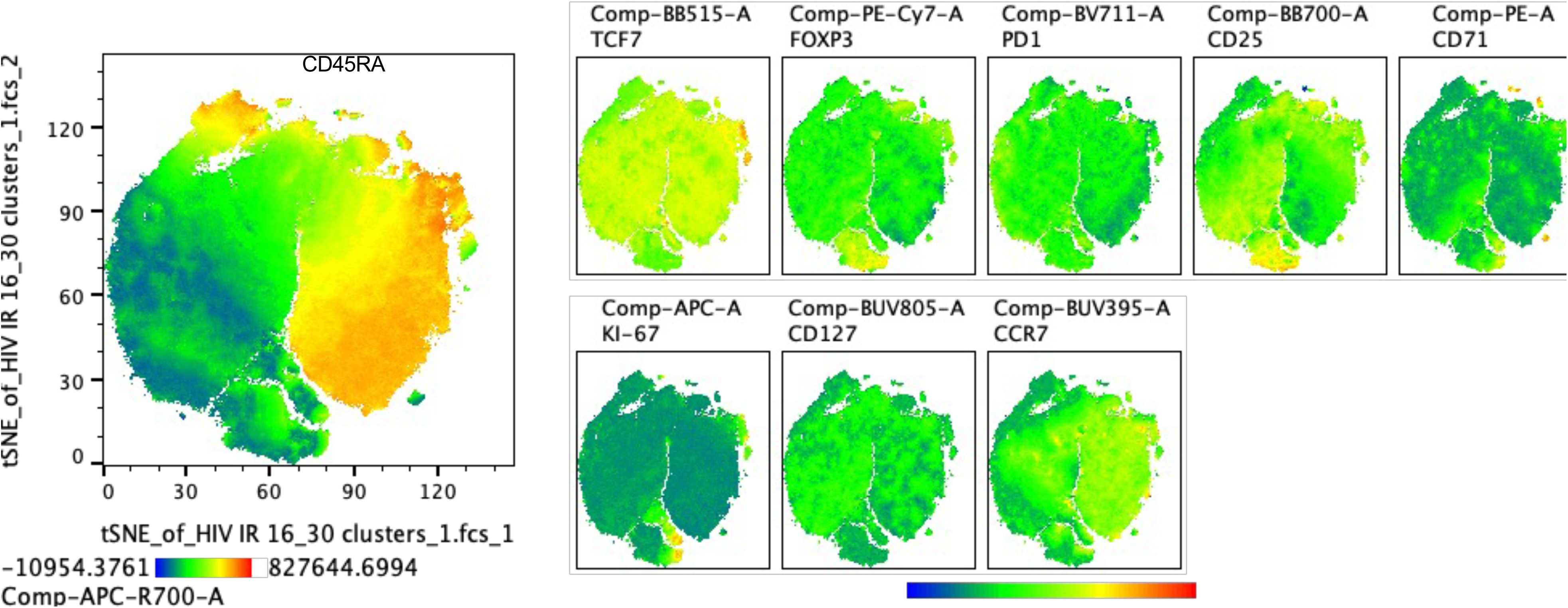
Marker distribution across CD4⁺ T-cell clusters used to define populations in Figure 2C. t-SNE projection of CD4⁺ T-cells from PLWH showing single-marker expression intensities across 30 FlowSOM-defined clusters. Each plot represents the expression of the indicated surface or intracellular marker (CD45RA, TCF7, FOXP3, PD-1, CD25, CD71, Ki-67, CD127, CCR7) across the CD4⁺ T-cell landscape. Expression intensities are color-coded from low (blue) to high (red). These marker distributions were used to assign phenotypic identity to CD4⁺ T-cell subsets shown in **Figure 2C**.

**Supplementary Figure 3.**
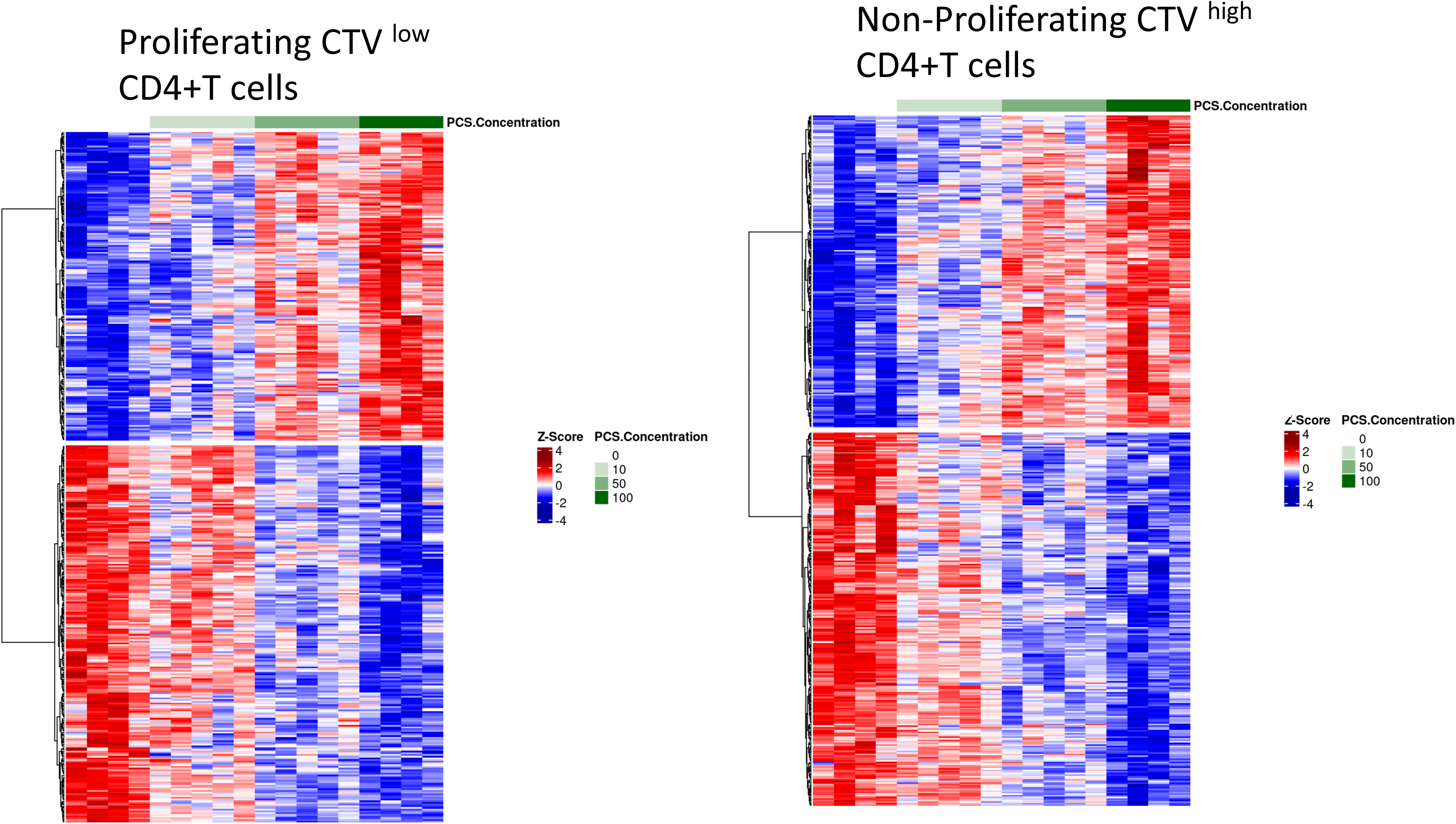
Transcriptomic response to PCS in proliferating and non-proliferating CD4⁺ T-cells. Heatmaps show hierarchical clustering of gene expression profiles (Z-score normalized) across PCS doses for proliferating (left) and non-proliferating (right) CD4⁺ T-cells. Transcriptional changes across both subsets showed broadly similar PCS-induced expression patterns. Based on this similarity, downstream transcriptomic and proteomic analyses were focused on the proliferating (CTV^low^) CD4⁺ T-cell subset.

**Supplementary Figure 4.**
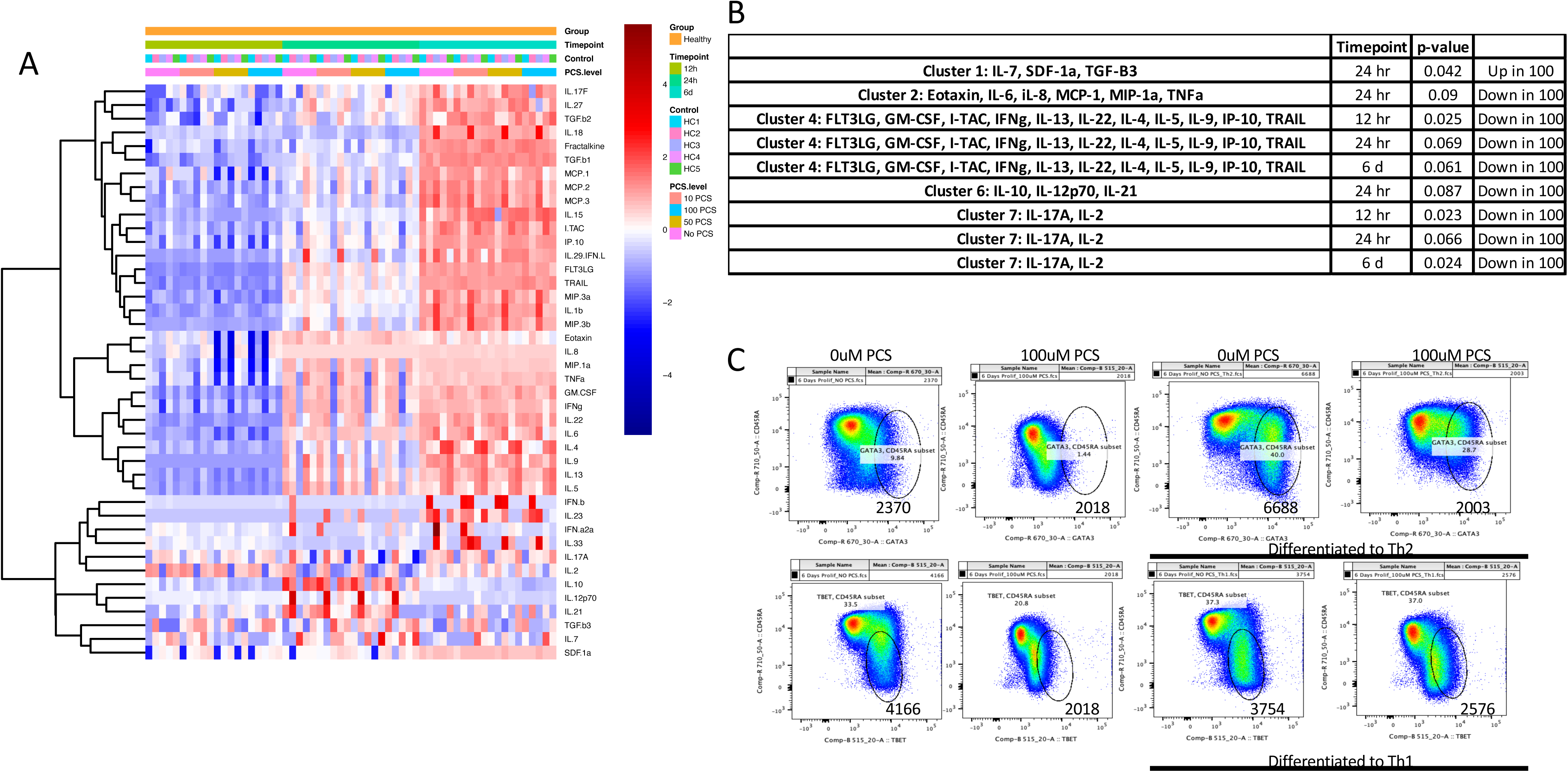
PCS modulates cytokine production and impairs Th1/Th2 polarization in CD4⁺ T-cells. **(A)** Heatmap of 40 cytokines measured in cell culture supernatants by multiplex detection system (MDS) from 5 healthy donors (HC1–HC5). PBMCs were stimulated with anti-CD3/CD28 and cultured in the presence of increasing concentrations of PCS (0, 10, 50, or 100 μM) for 12 hours, 24 hours, or 6 days. Cytokines were hierarchically clustered based on expression patterns. **(B)** Cluster analysis of cytokine profiles identified distinct PCS-responsive modules. Cluster 1 (IL-7, SDF-1a, TGF-β3) was upregulated at 24 hours with 100 μM PCS. In contrast, clusters 2, 4, 6, and 7, which include proinflammatory and T helper–associated cytokines (e.g., IL-6, TNF-α, IL-17A, IL-4, IL-5, IL-13), were significantly downregulated by PCS at multiple timepoints. Statistical significance determined by paired t tests. **(C)** PBMCs were stimulated with anti-CD3/CD28 for 6 days in the absence or presence of 100 μM PCS under non-polarizing (top left two panels), Th2-polarizing (top right two panels), or Th1-polarizing (bottom four panels) conditions. Cells were gated on CD4⁺ T-cells and analyzed for the expression of transcription factors GATA3 (top panels) and T-bet (bottom panels). The percentage of positive cells and the corresponding mean fluorescence intensity (MFI) values are shown on each plot. Data in **(C)** are representative of three independent experiments.

## Author contributions

SAY conceived and wrote the manuscript. ACDS, LK, JW, ALR, SK, SW, LDB, DEG, JAT performed experiments. ACS and SK performed scRNA-seq, AT analyzed scRNA-seq, JAT performed bulk RNAseq with ACDS, DEG and SW performed proteomics. VCM enrolled participants and obtained samples, JW, DJ and LK performed and analyzed metabolomics.

## Acknowledgments

This work was supported by the NIH grant R01AG076373 (SAY). VCM has received funding support from the Emory Center for AIDS Research (P30AI050409) for work related to this manuscript. We appreciate the assistance of the CFAR staff, Divine McCaslin, Ana Moldoveanu, and Shanil Fuller, and all the participants involved in this project.

## Disclaimers

VCM has received investigator-initiated research grants (to the institution) and research support from Eli Lilly, Bayer, Gilead Sciences, Merck, and ViiV.

## Declaration of generative AI and AI-assisted technologies in the writing process

During the preparation of this work, the authors used ChatGPT to improve the language and readability of the manuscript. After using this tool/service, the authors reviewed and edited the content as needed and take full responsibility for the content of the published article.

## Methods

### Flow cytometry

The panels used included the LIVE/DEAD™ Fixable Near-IR Dead Cell Stain Kit (catalog 501121531, Invitrogen, Waltham, MA, USA), anti-CD3 BUV737 (clone UCHT1, catalog 612750, BD Biosciences, Ashland, OR, USA), anti-CD4 BV605 (clone OKT4, catalog 317438, BioLegend, San Diego, CA, USA), anti-CCR7 BUV395 (clone 2-L1-A, catalog 569108, BD Biosciences, Franklin Lakes, NJ, USA), anti-CD45RA R718 (clone HI100, catalog 567073, BD Biosciences), anti-CD127 BUV805 (clone HIL-7R-M21, catalog 748486, BD Biosciences), anti-CD25 BB700 (clone M-A251, catalog 566447, BD Biosciences), anti-p21 Alexa Fluor™ 488 (clone EPR362, catalog ab282187, Abcam, Cambridge, UK), anti-p16INK4a PE (clone EPR1473, catalog ab209579, Abcam). Cells were fixed and permeabilized using the eBioscience FOXP3 / Transcription Factor Staining Buffer Set (catalog 00-5523-00, Invitrogen).

To perform the in vitro cell characterization assay for the Notch1 and Notch2 panel, cells from healthy donors were stimulated for 6 days with 200 ng of Purified NA/LE Anti-Human CD3 (clone UCHT1, catalog 555329, BD Biosciences) and Purified NA/LE Anti-Human CD28 (clone CD28.2, catalog 567117, BD Biosciences) in RPMI media (catalog 10-040-CV, Corning, Corning, NY, USA) containing 2% human serum (catalog 46026, Innovative Research, Novi, MI, USA) and 1% penicillin and streptomycin (catalog 30-002-CI, Corning). Cells were exposed to different concentrations of p-cresol sulfate (catalog A8895, APExBio, Houston, TX, USA): 10 μM, 50 μM, and 100 μM. The panel used included LIVE/DEAD™ Fixable Near-IR Dead Cell Stain Kit (catalog 501121531, Invitrogen), anti-CD3 BUV737 (clone UCHT1, catalog 612750, BD Biosciences), anti-CD4 BV605 (clone OKT4, catalog 317438, BioLegend), anti-CCR7 BUV395 (clone 2-L1-A, catalog 569108, BD Biosciences), anti-CD45RA R718 (clone HI100, catalog 567073, BD Biosciences), anti-Notch1 AF-647 (clone MHN1-519, catalog 564782, BD Biosciences), anti-Notch2 RB780 (clone MHN2-25, catalog 756225, BD Biosciences), anti-TCF-7/TCF-1 BV421 (clone S33-966, catalog 566692, BD Biosciences), anti-Beta-Catenin AF-488 (clone 14/Beta-Catenin, catalog 562505, BD Biosciences). P16-PE (clone EPR1473, catalog AB209579, Abcam), p21-AF488, clone EPR362, catalog AB282187, Abcam), AhR-PE-CY7 (clone FF3399, catalog 25-9854-42, eBioscience), CYP1B1-APC (polyclonal, catalogue NBP2-97885APC, Novus Biologicals). RB780 IgG2a kappa Isotype Control, (clone G155-178, catalog 568740, BD Biosciences), AF-488 IgG1 kappa Isotype Control (clone MOPC-21, catalog 557721, BD Biosciences), and BV421 IgG1 kappa Isotype Control (clone MOPC-21, catalog 569394, BD Biosciences).

### Th1/Th2/Treg Differentiation

To study the impact of PCS on CD4 differentiation, PBMCs from healthy donors were used for CD4 naïve T cell isolation – EasySep™ Human Naïve CD4+T-Cell Isolation Kit II (catalog 17555, STEMCELL Technologies, Vancouver, CA). Then, 500,000 CD4 naive cells were stimulated for 6 days with T Cell TransAct™, human (catalog 130-128-758, Miltenyi Biotec, Gaithersburg, MD, USA), at 1/3 of the recommended concentration in a 96-well U-bottom plate with AIM V™ Medium (catalog 12055091, Gibco, Waltham, MA, USA) and Nu-Serum™ IV Growth Medium Supplement (catalog 355504, Corning) and cultured in the presence of subset-specific cytokines in one final volume of 200μl. During this period, cells were also exposed to different PCS concentrations. For Treg differentiation naïve cells were stimulated with TransAct for 6 days in the presence of gradient PCS concentration. For Th1 differentiation, cells were stimulated with IL-12 (catalog 200-12H, PeproTech, Waltham, MA, USA) at 10 ng/mL, and anti-IL-4 antibody (catalog 130-095-753, Miltenyi Biotec) at 2.5 µg/mL. For Th2 differentiation, IL-4 (catalog 200-04, PeproTech) was used at 10 ng/mL, and anti-IFNγ antibody (catalog 130-095-743, Miltenyi Biotec) was used at 1 µg/mL. After 6 days, the panel used for Th1/Th2 differentiation included Live/Dead Fixable Near-IR (catalog 501121531, Invitrogen), anti-CD3 BUV737 (clone UCHT1, catalog 612750, BD Biosciences), anti-CD4 BV605 (clone OKT4, catalog 317438, BioLegend), anti-CCR7 BUV395 (clone, 2-L1-A catalog 569108, BD Biosciences), anti-CD45RA R718 (clone HI100, catalog 567073, BD Biosciences), anti-T-bet AF-488 (clone 4B10, catalog 568177, BD Biosciences), anti-Eomes PE (clone X4-83, catalog 566749, BD Biosciences), anti-GATA3 AF-647 (clone L50-823, catalog 560068, BD Biosciences), and TCF-7/TCF-1 BV421 (clone S33-966, catalog 566692, BD Biosciences). All cells were acquired using the BD FACSymphony A5 flow cytometer and analyzed with FlowJo™ v10.10.0 (BD Biosciences).

### Cytokine Profiling (MSD) Technology

Cytokine expression profiles were evaluated using a U-PLEX assay (Meso Scale MULTI-ARRAY^®^ Technology) available from Meso Scale Diagnostics (Rockville, MD, USA). This technology allows the evaluation of multiplexed biomarkers using custom-made U-PLEX sandwich antibodies with a SULFO-TAG conjugated antibody and next-generation electrochemiluminescence (ECL) detection. Pro-inflammatory and anti-inflammatory cytokines were analyzed, taking advantage of the platform’s broad dynamic range of detection (up to four logs, from 0.10 pg/mL to 1000 pg/mL). Combined with biotinylated antibodies plus the assigned linker and the SULFO-TAG conjugated detection antibody, the assay required 25 μL of plasma from each donor. Using a 10-spot U-PLEX plate, samples were read by the MESO QuickPlex SQ 120 to detect electrochemiluminescence specific to each of the ten spots per well. Results were reported in pg/mL and extrapolated from the standard curve for each specific analyte.

### Single-cell RNA-Seq

Frozen PBMCs were thawed, and total CD4^+^T-cells were enriched using the EasySep™ Human CD4^+^T-Cell Enrichment Kit (StemCell Technologies, Vancouver, CA), following the manufacturer’s instructions. Cells were washed with PBS containing BSA 0.04%, and viability was assessed by staining the cells with Trypan blue. Cells were resuspended at the desired concentration to target 10,000 cells per sample. Single-cell suspensions were prepared using the Chromium Next GEM Single Cell 5’ Reagent Kits v2 (10x Genomics, Pleasanton, CA) and partitioned into Gel Beads-in-emulsion (GEMs) using the 10x Chromium Controller and Next GEM Chip K. GEMs were visually inspected, and only samples with an opaque and uniform aspect were used for library preparation. Gene Expression (GEX) libraries were generated from the cDNA. Representative traces and quantitation of libraries were determined using Bioanalyzer High Sensitivity DNA Analysis (Agilent, Santa Clara, CA). GEX libraries were sequenced on Illumina NovaSeq S4 system, using 10x’s recommended parameters. The 10x Barcodes in the library were used to associate individual reads back to the individual partitions, and thereby, to each single cell.

### Bulk RNA-seq

Cells from healthy donors were labeled with CellTrace™ Violet (catalog C34557, Invitrogen) and stimulated for 6 days with 200 ng of purified NA/LE anti-Human CD3 (clone UCHT1, catalog 555329, BD Biosciences) and purified NA/LE anti-Human CD28 (clone CD28.2) in RPMI media (catalog 10-040-CV, Corning) containing 2% human serum (catalog 46026, Innovative Research) and 1% penicillin and streptomycin (catalog 30-002-CI, Corning). Cells were exposed to different PCS concentrations (catalog A8895, APExBIO): 10 μM, 50 μM, and 100 μM. After 6 days of stimulation and PCS exposure, CD4 proliferated cells were sorted by BD FACSAria™ II (BD Biosciences), followed by RNA isolation using the RNeasy Kit (catalog 74104, QIAGEN, Venlo, NL) and library preparation. Library preparation was performed according to the manufacturer’s instructions using the Illumina^®^ Stranded Total RNA Prep with Ribo-Zero Plus Kit (catalog 20040529, Illumina, San Diego, CA, USA), and IDT for Illumina^®^ RNA UD Indexes Set A, Ligation (96 Indexes, 96 Samples, catalog 20040553). cDNA library quality was assessed using an Agilent High-Sensitivity DNA Kit (catalog 5067-4626, Agilent Technologies, Santa Clara, CA, USA), yielding a broad peak between 300-800 base pairs. Sequencing was performed by the Emory Non-primate Research Center Genomics Core using the Illumina NovaSeq S4 platform and the NovaSeq S4 200-cycle Kit (catalog 20028313, Illumina) to achieve high read depth. Raw reads were assessed using FastQC and aligned to the GRCh38 genome using the STAR aligner. Only samples meeting quality control thresholds were included in further analysis, ensuring alignment with expression patterns from traditional bulk RNA sequencing studies.

### Proteomics Sample Preparation

Cells were washed in PBS and frozen before proteomics analysis. Frozen cells were resuspended in cold PBS with cOmplete Mini EDTA-free Protease Inhibitor Cocktail (catalog 11836170001, Roche, Basel, CH) and sonicated using a Bioruptor (15s on / 15s off, 10 cycles x 2; Hologic Diagenode, Liège, BE). Protein was quantified by Pierce BCA Protein Assay (catalog 23225, Thermo Fisher Scientific, Waltham, MA, USA). Samples were diluted 1:10 into RAD buffer: 100 mM Tris (pH 8.8) containing 1 % SDC, 10 mM TCEP, and 40 mM CAA. Samples were then boiled at 95 °C for 10 minutes, then cooled to RT. Proteins were digested overnight at 37 °C with Trypsin/Lys-C Mix (1:40 w/w; catalog V507A, Promega Corp., Madison, WI, USA). Peptides were acidified with formic acid and centrifuged to pellet precipitated deoxycholate. Peptides were cleaned up on Oasis 2 mg HLB plates (catalog 186001828ba, Waters Corp., Milford, MA, USA) and quantified using a Pierce Quantitative Colorimetric Peptide Assay (catalog 23275, Thermo Fisher Scientific) according to the manufacturer’s instructions then dried in a SpeedVac Vacuum Concentrator (Thermo Fisher Scientific).

### Proteomics LC-MS/MS

Samples were resuspended in buffer A (0.1% FA in water) and 200 ng per sample was injected in a Bruker NanoElute coupled to a timsTOF Pro2 mass spectrometer (Bruker Daltonics, Billerica, MA, USA). The peptides were loaded on a 25 cm Aurora Elite CSI column (IonOpticks, Collingwood, AU). The mass-spectrometer operated in positive polarity for data collection using data-independent acquisition (diaPASEF) mode.

### Proteomics Data Analysis

Protein identification and quantification were performed using Spectronaut in library-free mode using the Homo sapiens SwissProt database with default settings. Modifications were defined as follows: Carbamidomethylation (C) as a fixed modification, and Acetyl (Protein N-term) and Oxidation (M) as variable modifications.

### Metabolic Analysis

Untargeted and targeted metabolic analysis of isolated CD4^+^T-cells and plasma from PLWH donors was performed at the Clinical Biomarkers Laboratory in the Emory University School of Medicine.

### Untargeted High-Resolution Metabolomics

Cells were extracted with 200 μL of ice-cold 80:20 acetonitrile:water plus internal standards and incubated on ice for 10 mins. Plasma was extracted with a 2:1 ratio using ice-cold acetonitrile spiked with internal standards and incubated on ice for 30 min. All samples then underwent centrifugation at 14,000 rpm for 10 min at 4 °C. Supernatants were transferred to autosampler vials and stored at 4 °C for analysis on the same day. Samples were randomized and analyzed in triplicate on a dual column liquid chromatography system (Thermo Ultimate3000 UHPLC, Thermo Fisher Scientific) connected to a high-resolution mass spectrometer (Thermo Q Exactive HF, Thermo Fisher Scientific). The two analytical platforms consisted of a Waters Xbridge BEH Amide XP HILIC column (2.1 mm x 50 mm, 2.6 μm particle size) coupled to positive electrospray ionization (ESI+) and a Higgins Targa C18 column (2.1 mm x 50 mm, 3 μm, Thermo Fisher Scientific) coupled to negative electrospray ionization (ESI-). The mobile phases included LCMS grade water (A), LCMS grade acetonitrile (B), and 2% formic acid in water (C), and 10 mM ammonium acetate in water (D). For the HILIC method, a buffer ratio of 22.5% A, 75% B, and 2.5% C was held for 1.5-min before ascending to 75% A, 22.5% B, and 2.5% C for 4 min and ending with a 1-min gradient hold. For the C18 gradient, an initial buffer ratio of 60% A, 35% B, and 5% D was held for 1-min before ascending to 0% A, 95% B, and 5% D over 3 min and ending with a 2-min gradient hold. Flow rates were set at 0.35 mL/min for the first min and 0.4 mL/min for the remaining 4 minutes. Column compartments were heated to 60 °C. Mass spectra were collected at 120k resolution in an 85-1,275 *m/z* scan window.

### Metabolomics Feature Extraction

Mass spectral feature extraction was performed with adaptive processing for LCMS (apLCMS) (v6.3.3) coupled to quality control processing by xMSanalyzer (v2.0.8). Features underwent peak detection, noise removal, alignment, and technical triplicates were median summarized. The resulting features had a corresponding retention time, *m/z*, and peak areas. False discovery rate (FDR) correction by Benjamini-Hochberg was applied using a 5% threshold. Annotation of features was performed with xMSannotator and the Human Metabolome Database as the reference library.

### Targeted Quantification of Selected Metabolites

For metabolite extraction, cells were treated as previously described(80). Quantification was performed by method of standard addition for PCS, PCG, PAG, and IAA. Cell extracts, standard curves, and metabolite standards were analyzed in triplicate on a Vanquish Duo UHPLC (Thermo Fisher Scientific) coupled to an Orbitrap ID-X mass spectrometer (Thermo Fisher Scientific) with similar HILIC/ESI+ and C18/ESI- methods as previously described. Mass spectra were collected at 60k resolution in a 170-310 *m/z* scan window. Targeted *m/z’*s then underwent MS^2^ ion dissociation at 35 % normalized collision energy (HCD). MS^2^ scans were acquired at 15k resolution. Peak area determination and MS^2^ spectra evaluation were performed with Thermo xCalibur (v4.2) QualBrowser.

### HIV Provirus Detection

PBMCs from our cohort were sent to Accelevir Diagnostics (Baltimore, MD, USA) for IPDA™ (Intact Proviral DNA Assay) analysis.

### Study Participants and Approval

Fifty PLWH, all receiving antiretroviral therapy (ART) having undetectable plasma HIV-1 RNA viral loads, were identified from the Emory Center for AIDS Research (CFAR) HIV Disease Registry and recruited from the Ponce Center, Grady Health System to participate in this study. All interested and eligible participants provided written informed consent under an approved protocol approved in accordance with the ethical principles of the Declaration of Helsinki.The cohort had a median age of 59 years (range: 40–70), a median CD4^+^T-cell count of 735 cells/μL (range: 478–1453), and a median duration on ART of 10.15 years (range: 0.2–25.23 years). From this group, 26 participants were randomly selected to undergo both untargeted and targeted metabolomic analyses.

