## Supplemental Figures 1-4 for "Microbiome-Derived Metabolites Shape CD4⁺ T-Cell Differentiation and Immune Aging in Chronic HIV-1 Infection"

**Supplementary Figure 1**

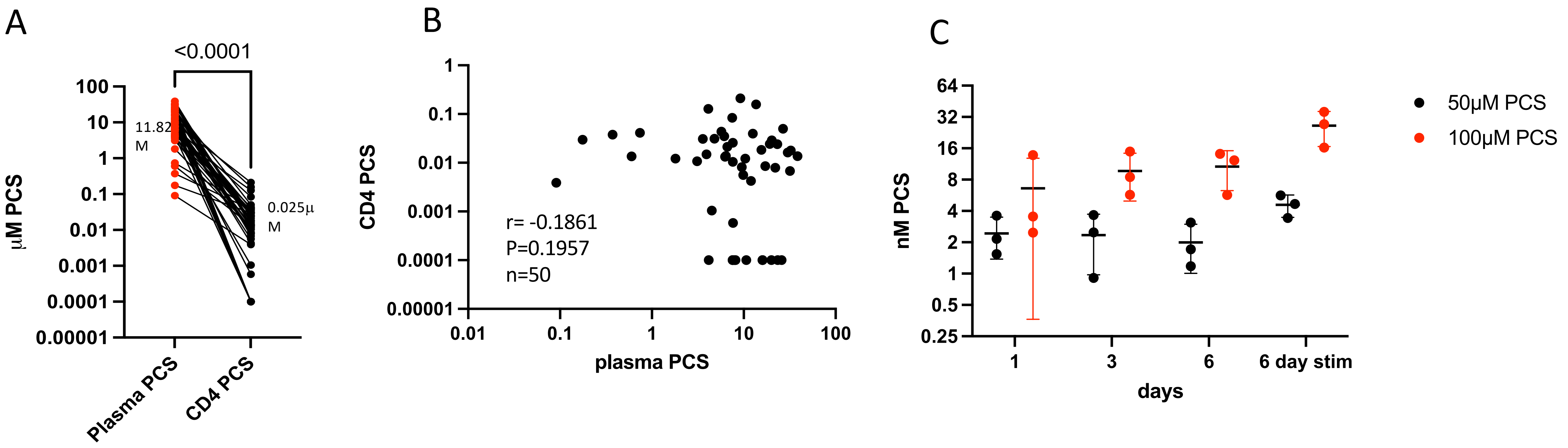

**Supplemental Figure 2**

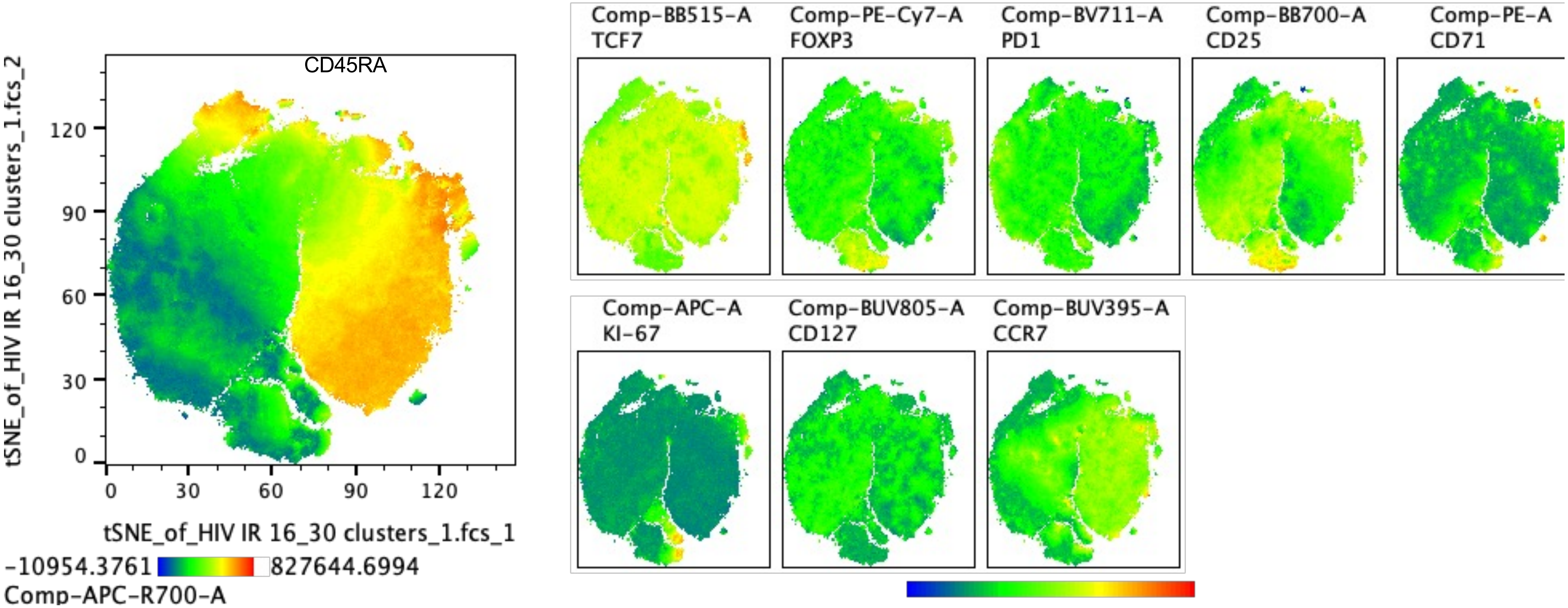

### Supplemental Figure 3

Proliferating CTV $^{low}$ CD4+T cells

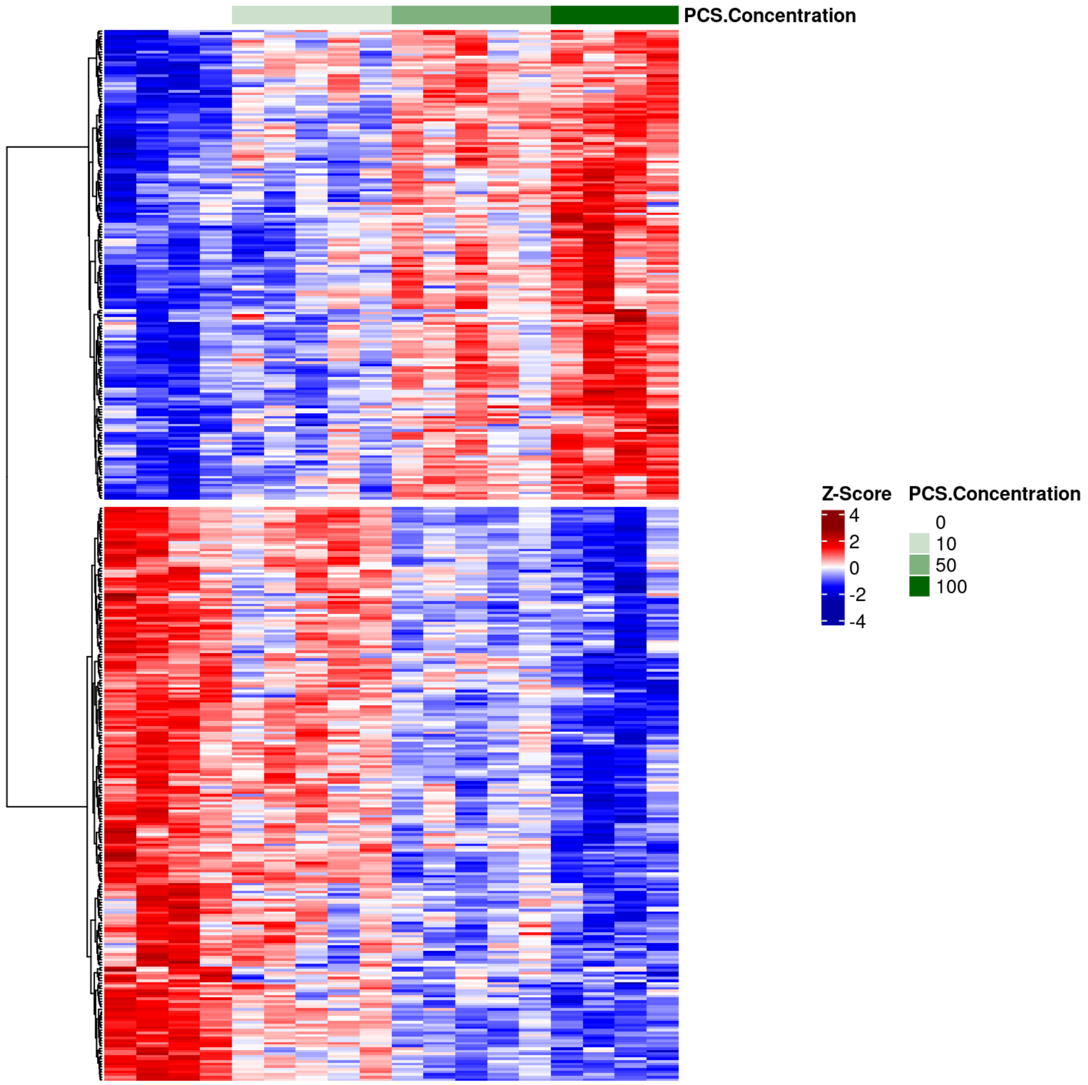


Non-Proliferating CTV $^{high}$ CD4+T cells

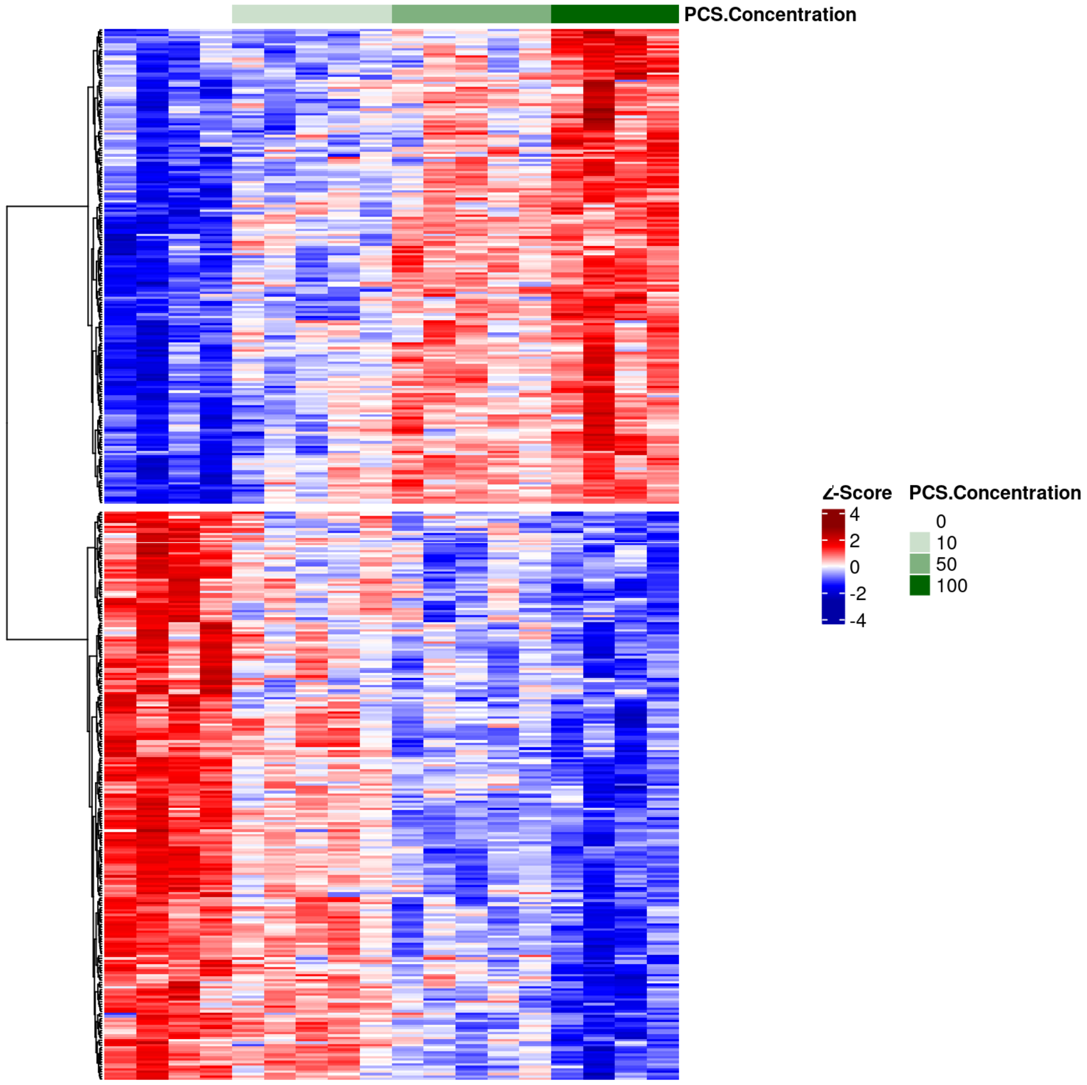

### Supplementary Figure 4

A

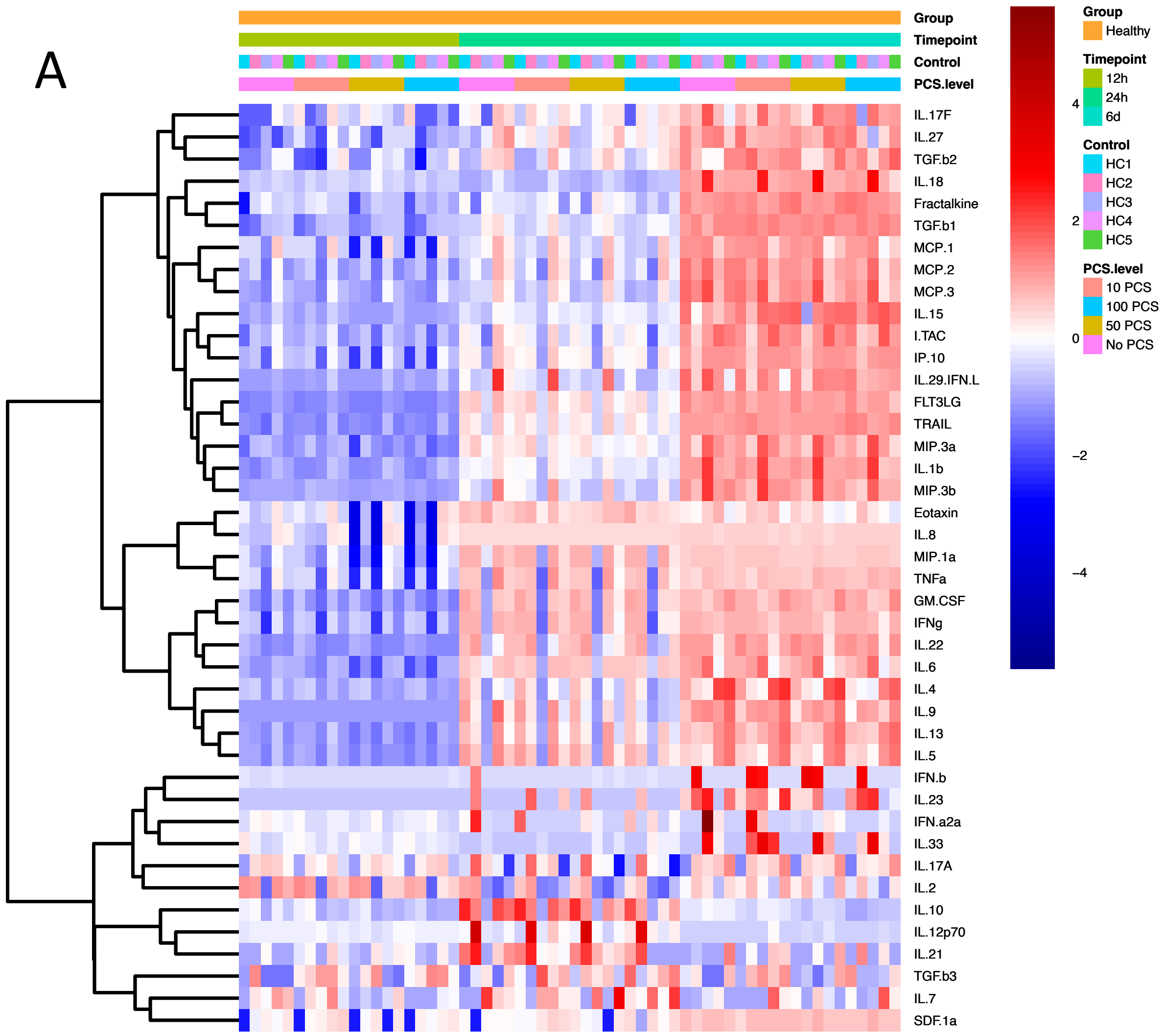


B

| | Timepoint | p-value | |
|---|---|---|---|
| Cluster 1: IL-7, SDF-1a, TGF-B3 | 24 hr | 0.042 | Up in 100 |
| Cluster 2: Eotaxin, iL-6, iL-8, MCP-1, MIP-1a, TNFa | 24 hr | 0.09 | Down in 100 |
| Cluster 4: FLT3LG, GM-CSF, I-TAC, IFNg, IL-13, IL-22, IL-4, IL-5, IL-9, IP-10, TRAIL | 12 hr | 0.025 | Down in 100 |
| Cluster 4: FLT3LG, GM-CSF, I-TAC, IFNg, IL-13, IL-22, IL-4, IL-5, IL-9, IP-10, TRAIL | 24 hr | 0.069 | Down in 100 |
| Cluster 4: FLT3LG, GM-CSF, I-TAC, IFNg, IL-13, IL-22, IL-4, IL-5, IL-9, IP-10, TRAIL | 6 d | 0.061 | Down in 100 |
| Cluster 6: IL-10, IL-12p70, IL-21 | 24 hr | 0.087 | Down in 100 |
| Cluster 7: IL-17A, IL-2 | 12 hr | 0.023 | Down in 100 |
| Cluster 7: IL-17A, IL-2 | 24 hr | 0.066 | Down in 100 |
| Cluster 7: IL-17A, IL-2 | 6 d | 0.024 | Down in 100 |

C

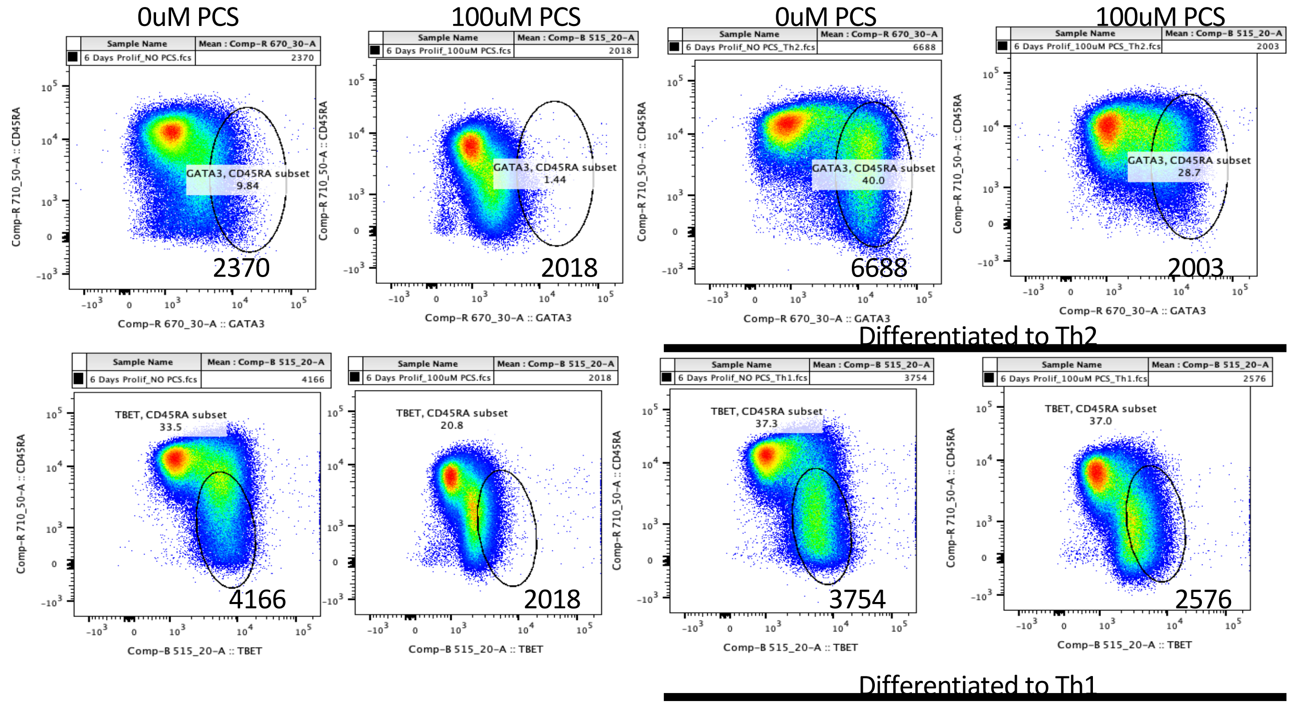
